# Water for sterol: an unusual mechanism of sterol egress from a StARkin domain

**DOI:** 10.1101/623777

**Authors:** George Khelashvili, Kalpana Pandey, Neha Chauhan, David Eliezer, Anant K. Menon

**Affiliations:** Department of Physiology and Biophysics, Weill Cornell Medical College, 1300 York Ave, New York, NY 10065, USA; Institute for Computational Biomedicine, Weill Cornell Medical College, 1300 York Ave, New York, NY 10065, USA; Department of Biochemistry, Weill Cornell Medical College, 1300 York Ave, New York, NY 10065, USA

## Abstract

Previously we identified a new family of endoplasmic reticulum membrane proteins that possess sterol-binding StARkin domains (Gatta et al. eLife 2015). These Lam/GramD1 proteins are implicated in intracellular sterol homeostasis, a function that requires them to be able to bind sterols. Here we show how these proteins exchange sterol molecules with membranes. An aperture at one end of the StARkin domain enables sterol to enter/exit the binding pocket. Strikingly, the wall of the pocket is fractured along its length, exposing bound sterol to solvent. We considered whether hydration of the pocket could mediate sterol entry/exit. Large-scale atomistic molecular dynamics simulations reveal that sterol egress involves widening of the fracture, penetration of water into the cavity and consequent destabilization of the bound sterol. The simulations also identify polar residues along the fracture that are important for sterol release. Their replacement with alanine affects the ability of the StARkin domain to bind sterol, catalyze inter-vesicular sterol exchange and alleviate the nystatin-sensitivity of *lam2Δ* yeast cells. These data suggest an unprecedented, water-controlled mechanism of sterol acquisition and discharge from a StARkin domain.

## Introduction

Cholesterol, the ‘central lipid of mammalian cells’ (1), is the most abundant molecular component of the mammalian plasma membrane (PM) where it represents one out of every 2-3 lipids (1,2). Like many membrane lipids, it is synthesized in the endoplasmic reticulum (ER) and transported to the PM by non-vesicular mechanisms that make use of lipid transport proteins (3,4). These proteins operate as molecular ferries, achieving lipid exchange between membranes by reversibly extracting a lipid from the cytoplasmic leaflet of one membrane bilayer, encapsulating it within a binding pocket for transfer through the cytoplasm, and depositing it in the cytoplasmic leaflet of another membrane. Proteins with steroidogenic acute regulatory protein related lipid transfer (StART) domains constitute a major family of intracellular lipid transport proteins - the StARkin superfamily - implicated in moving glycerophospholipids, ceramide and sterol between cellular membranes (5,6). Whereas these proteins are generally soluble and able to diffuse freely through the cytoplasm, a new family of ER membrane proteins with StARkin domains was recently identified, including six members (Lam1-Lam6) in the budding yeast *Saccharomyces cerevisiae* and three members (GramD1a-GramD1c) in mammals (7-10). Members of this new sub-family have one or two StARkin domains that bind sterols and catalyze sterol exchange between populations of vesicles *in vitro* (7,9,11-14). Lam1-Lam4 localize to ER-PM contact sites in yeast (7,15) where they play a role in sterol homeostasis. Thus, yeast cells lacking one or more of these proteins are hypersensitive to the sterol-binding, polyene antibiotics amphotericin and nystatin, implying alterations in PM sterol content and/or organization (7,16). Furthermore, they esterify exogenously supplied sterols up to 3-fold more slowly than wild-type cells, indicative of a delay in some aspect of PM-ER sterol transport (7,16). A sterol homeostatic role has also been suggested for mouse GramD1b protein which is highly expressed in steroidogenic organs. Thus, adrenal glands from a GramD1b knockout mouse are devoid of lipid droplets and show a severe reduction in cholesteryl ester content (14).

We recently reported crystal structures of the second StARkin domain of Lam4 (here termed Lam4S2) in *apo*- and sterol-bound states (11). The protein has an overall α/β helix-grip fold that forms a capacious binding pocket into which the sterol appears to be admitted head-first, via an aperture at one end, such that its 3-β-hydroxyl head-group is stabilized by direct or water-mediated interactions with polar residues (***Figure 1A***). The surface of the protein near the entrance to the pocket is decorated with lysine residues, accounting for the enhanced ability of Lam4S2 to transfer sterol between anionic vesicles compared with neutral vesicles (11); the entryway itself is partially occluded by a flexible loop, termed Ω_1_ (***Figure 1A***), whose functional importance in the StARkin family is well-documented through mutagenesis studies (12,17,18). The structures of other Lam/GramD1 StARkin domains are similar (12-14), broadly resembling structures of other members of the StARkin superfamily except for one striking feature. The wall of the sterol binding cavity in all Lam/GramD1 StARkin domains is fractured along part of its length, exposing the sterol backbone to bulk solvent (***Figure 1-figure supplement 1***). We considered whether this unusual structural feature - henceforth termed ‘side-opening’ - might provide a mechanism to control the stability of sterol within the binding pocket. We posited that the ability to load sterol into the pocket, or discharge it from the pocket into the membrane, might be controlled by water permeation via the side-opening. We used a combination of large-scale atomistic molecular dynamics (MD) simulations and functional tests to explore this hypothesis. Analysis of extensive ensemble and umbrella sampling MD trajectories revealed that sterol egress from Lam4S2 is associated with widening of the side-entrance to the binding pocket, penetration of water molecules into the cavity and consequent destabilization of the bound sterol. The simulations identified several polar residues that line the side-entrance to the pocket and that appear to play a critical role in the initial steps of the release process. The functional importance of these residues was validated experimentally by showing that their replacement with alanine compromises the ability of Lam4S2 to rescue the nystatin-sensitivity of *lam2Δ* yeast cells and reduces the efficiency with which the purified protein is able to extract membrane-bound sterol and catalyze sterol exchange between populations of vesicles *in vitro*. These data suggest an unprecedented, water-controlled mechanism of sterol acquisition and discharge from a StARkin domain.

**Figure 1.**
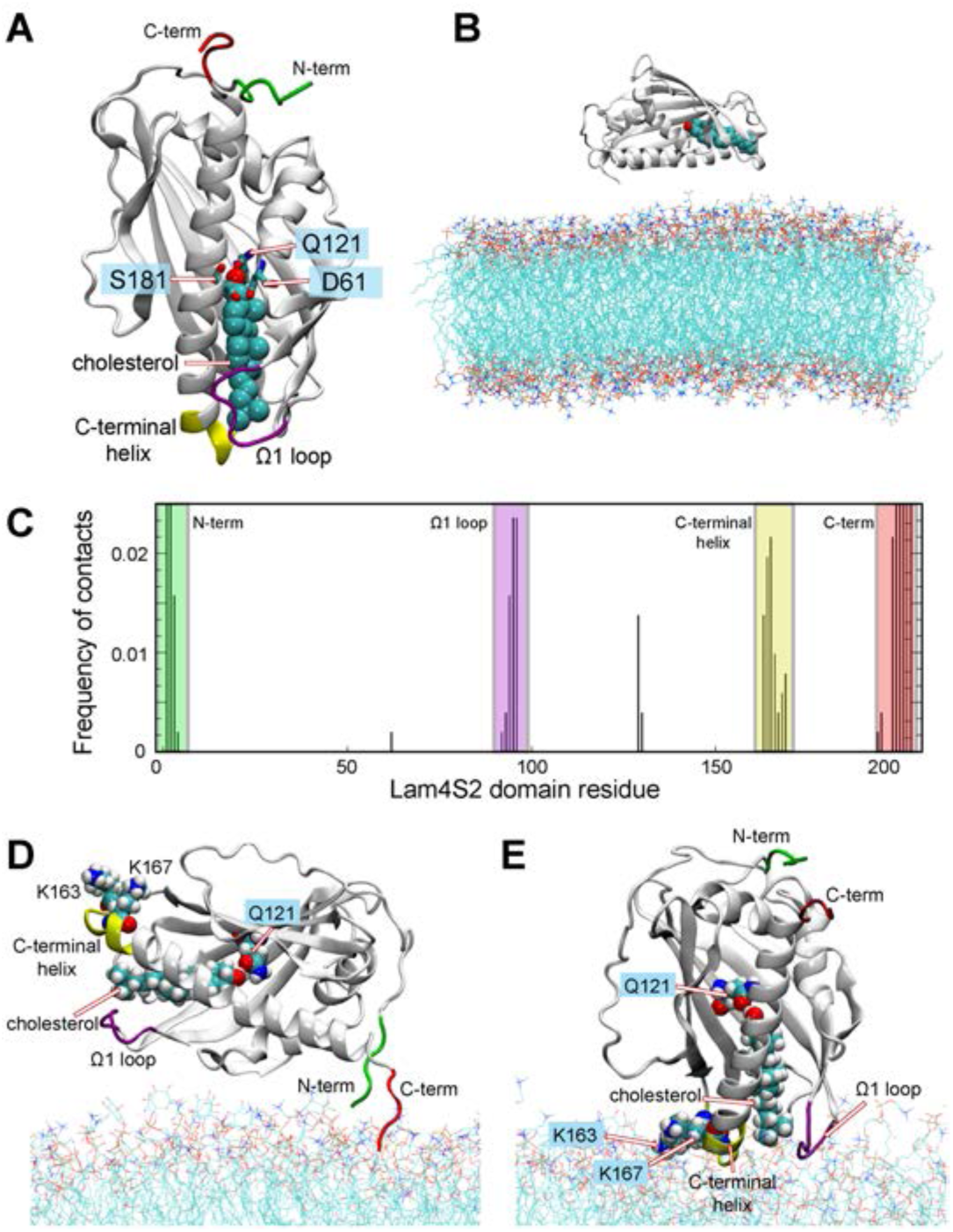
Different modes of Lam4S2-membrane association deduced from ‘Stage 1’ molecular dynamics simulations. (**A**) Structural elements of the Lam4S2 domain used to construct collective variables (CVs) for tICA analysis. Shown in licorice and labeled are residues Q121, S181, D61. Also highlighted are locations of the Ω_1_ loop (purple), the C-terminal helix (yellow), the N-terminus (green), and the C-terminus (red). Cholesterol is shown in space filling representation colored cyan except for the oxygen atom in red (**B**) Initial positioning of Lam4S2 (cartoon) near the membrane (lines). In this configuration, the distance between any atom of the protein and any atom of a lipid molecule was ≥10Å. The cholesterol molecule bound to Lam4S2 is shown in space-fill representation. The water box including solution ions are omitted for clarity. (**C**) For each residue of the Lam4S2, the fraction of trajectory frames from Stage 1 simulations during which the residue is in contact with the membrane is plotted. A residue was considered in contact with the bilayer if the z-coordinate of the C_α_ atom of this residue was within 1Å of the average z-position of the neighboring lipid phosphate atoms (identified as the ones located within 10Å of this C_α_ atom). The relevant protein segments are labeled and colored using the color-code used in panel A. (**D-E**) Two modes of Lam4S2-membrane association. The lipids in the membrane are shown as lines. The relevant protein segments are labeled and colored using the color-code used in panel A. For completeness, panels D-E also show in space-fill the protein-bound cholesterol molecule, and residues K163, K167, and Q121.

## Results

### Lam4S2 associates with the membrane via its Ω1 loop and C-terminal helix

We used atomistic MD simulations of Lam4S2 to explore the impact of the unique lateral opening in Lam/GramD1 StARkin domains on the stability of bound sterol and its ability to exit the binding pocket. Cholesterol-bound Lam4S2 (***Figure 1A***) was placed in the vicinity of a membrane bilayer (***Figure 1B***) having the composition of anionic “Acceptor” liposomes used for *in vitro* sterol transport assays (11), and its spontaneous binding to the membrane surface was monitored via ensemble MD simulations carried out in 10 statistically independent replicates (Stage 1 ensemble simulations, 3.2 µs cumulative time). The simulations showed two modes by which Lam4S2 associated with the membrane. In one mode, the protein interacted with lipid headgroups via its N- and C-termini (green and red; ***Figure 1C, D***). As these regions are linked to adjacent parts of the protein chain in full-length Lam4, i.e. the S1 domain and the transmembrane helix, respectively, this mode of association is likely a non-physiological artifact of simulating the isolated Lam4S2 domain and was not pursued further. In the second mode, Lam4S2 engaged with the membrane via its Ω1 loop (purple; ***Figure 1C, E***) and its C-terminal helix (yellow; ***Figure 1C, E***). The former is a functionally important feature of all StARkin domains (12,17,18), whereas the latter harbors cationic residues (e.g., K163, K167) which have been implicated in the *in vitro* sterol transfer activity of Lam4S2 (11,17) and the protein StARD4 (18). In this binding mode, cholesterol is oriented orthogonally to the plane of membrane, with its 3-β-OH group engaging Q121 in the binding pocket of Lam4S2 and its iso-octyl tail facing the membrane (***Figure 1E***). We hypothesized that in this position the protein is primed to release sterol into the membrane and proceeded to test this premise by enhancing the sampling of this mode of Lam4S2-membrane interaction. To this end, we initiated a new set of 100 independent MD simulations (with random starting velocities) from 10 conformations of the system in which the Ω1 loop and the C-terminal helix were simultaneously engaged with lipids (Stage 2 ensemble simulations, 37.5 µs cumulative time). As described next, these trajectories revealed detailed mechanistic steps leading to spontaneous release of the protein-bound cholesterol into the membrane.

### Mechanistic steps of cholesterol transfer from Lam4S2 to the membrane

To facilitate analyses of the conformational dynamics of the membrane-bound Lam4S2-cholesterol complex in Stage 2 simulations, we used the time-structure based independent component analysis (tICA) approach to reduce the dimensionality of the system (see Methods). To this end, we considered a set of collective variables (CVs) to describe the dynamics of cholesterol and relevant segments of the protein (i.e. the Ω_1_ loop and the C-terminal helix) as well as to quantify solvent exposure of the sterol binding site (see Methods for details). All the trajectory frames from Stage 2 simulations were projected on the first two tICA vectors, which represented ∼90% of the total dynamics of the system (***Figure 2-figure supplement 2A***). The resulting 2D space (***Figure 2A***), was discretized for structural analyses into 100 microstates using the automated *k*-means clustering algorithm (***Figure 2-figure supplement 2B***). These microstates cover the conformational space of the system as the cholesterol molecule is transferred from the protein-bound state to the membrane.

**Figure 2:**
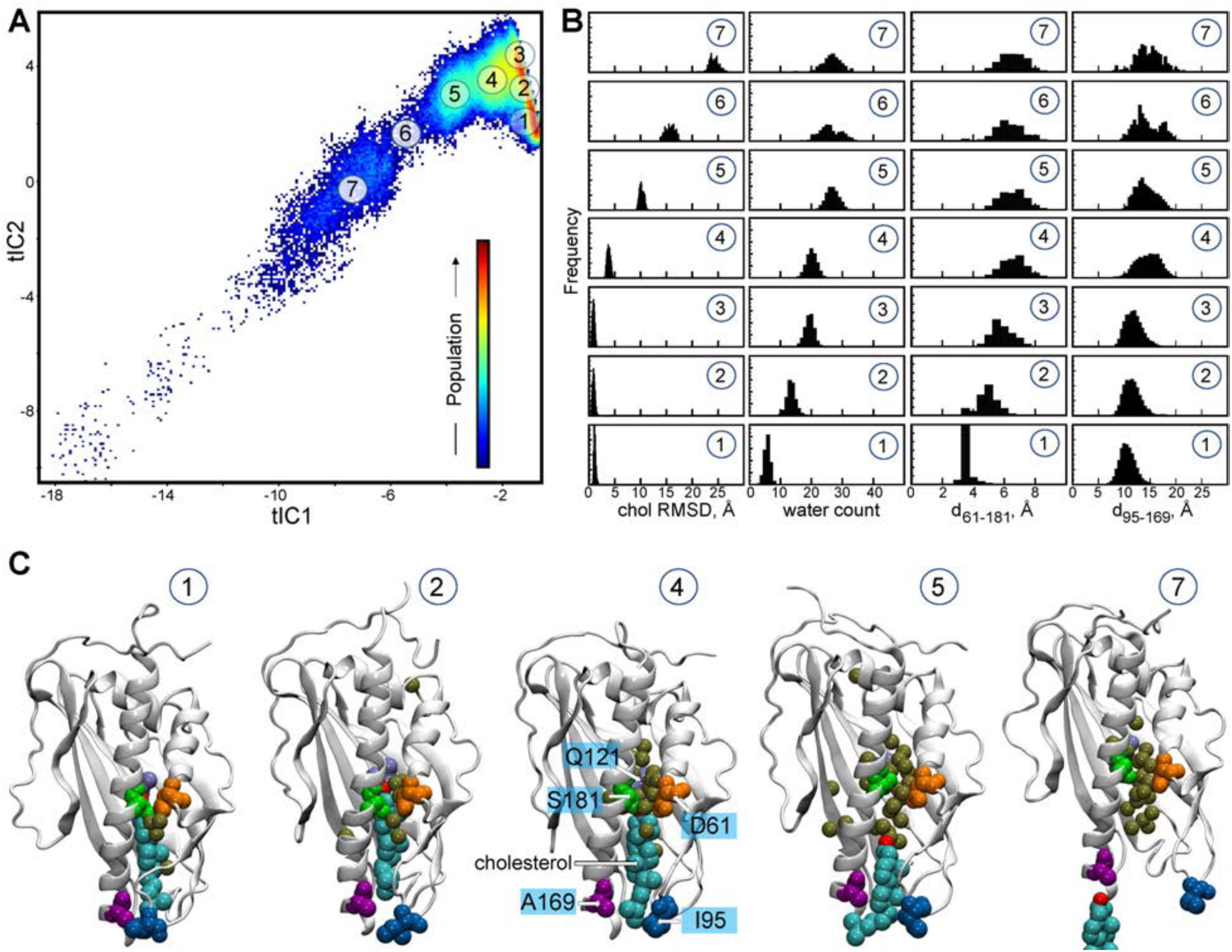
Mechanistic steps of cholesterol release from Lam4S2 revealed from tICA analysis. (**A**) 2-D landscape representing all the Stage 2 MD trajectories mapped with the tICA transformation in the space of the first two tICA eigenvectors (tIC 1 and tIC 2). The lighter shades (from red to light green to yellow) indicate the most populated regions of the 2D space (see the color bar). Microstates representing the most populated states in these simulations are indicated by the numbered circles (1-7) and represent various stages in the lipid translocation process. (**B**) Structural characteristics of the selected 7 microstates. The columns record the probability distributions of the cholesterol RMSD, number of water oxygens in the sterol binding pocket, and distances between residues 61 and 181 (d_61-181_) and 95 and 169 (d_95-169_). (**C**) Structural models representing selected microstates. In these snapshots, Lam4S2 is shown in cartoon, and cholesterol as well as selected protein residues (Q121, D61, S181, I95, A169) are shown in space fill (the residues are labeled in the snapshot of Microstate 4). Water oxygens in the sterol binding site are drawn as gold spheres.

Structural analyses of selected microstates on the tICA landscape (labeled 1-7 in ***Figure 2A***), characterized by relevant CVs (***Figure 2B***) and visualized in structural snapshots (***Figure 2C***) describe key mechanistic steps of the sterol release process. Microstate 1 represents an ensemble of states in which the sterol binding cavity is occluded from both the solvent and the membrane. Thus, in Microstate 1 conformations (***Figure 2B, C***), cholesterol is stably bound in the protein (“chol RMSD” histogram (bottom panel, ***Figure 2B***)), while the sterol binding pocket is dehydrated (“water count” histogram (bottom panel, ***Figure 2B***)) and sealed from the side by the side-chains of residues S181 and D61 that line the side-entrance to the pocket (d_61-181_ distance histogram (bottom panel, ***Figure 2B***)). In addition, the Ω_1_ loop is positioned close to the C-terminal helix so that the C_α_ atoms of residues I95 in the Ω_1_ loop and A169 in the C-terminal helix are within ∼10Å of each other (d_95-169_ distance histogram (bottom panel, ***Figure 2B***); see the middle structure in ***Figure 2C*** for the location of A169), therefore occluding the sterol binding pocket from below, i.e. the vantage point of the membrane. Indeed, as shown in ***Figure 2-figure supplement 3***, in the ensemble of conformations representing Microstate 1, cholesterol has essentially no contact with membrane lipids (“number of lipids” histogram (bottom panel, ***Figure 2-figure supplement 3B***)).

The first step in the sterol release process involves widening of the side-entrance to the sterol binding site enabled by gradual separation of the side-chains of residues D61 and S181. This structural change on the tICA landscape can be followed in the evolution of the system from Microstate 1 to Microstates 2 and 3 (see d_61-181_ distance histogram (***Figure 2B***)). Concomitant with the widening of the side-entrance, the level of hydration (water count) of the binding pocket progressively increases (***Figure 2B, C***).

Cholesterol remains stably bound throughout these initial events (cholesterol RMSD is unchanged in Microstates 1-3). However, the rising level of hydration in the binding site results in destabilization of the polar interactions between the 3-β-OH group of cholesterol and the side-chain of residue Q121 as cholesterol initiates its translocation towards the membrane. Indeed, as the system transitions from Microstate 3 to Microstate 4, the RMSD of the cholesterol molecule increases (***Figure 2B***). Correspondingly, the minimum distance between the cholesterol oxygen and residue Q121 increases by ∼4Å (compare d_chol-121_ for Microstates 1 and 4, ***Figure 2-figure supplement 3***). Notably, as the cholesterol molecule assumes this new position, the distance between the Ω_1_ loop and the C-terminal helix increases as seen in the broadening of the d_95-169_ histogram (***Figure 2B***), indicating initial opening of the sterol binding pocket towards the membrane.

Cholesterol egress then proceeds through Microstates 5-7 in which the sterol binding pocket remains open and solvated, while the Ω_1_ loop continues to sample conformations that position it relatively far from the C-terminal helix (***Figure 2B, C***). The cholesterol molecule leaves the binding pocket with its tail “down”, becoming gradually encapsulated by the hydrophobic chains of neighboring lipids until it fully embeds into the lipid membrane (“number of lipids” histogram (bottom panel, ***Figure 2-figure supplement 3B***) and corresponding structural snapshots (***Figure 2-figure supplement 3D***)). The process of translocation is complete when the system reaches Microstate 7. The remaining part of the tICA space (corresponding to lower tIC1 and tIC2 values, i.e. bottom left region of the 2D space in ***Figure 2A***), describes trajectory data in which Lam4S2 disengages from the membrane, after the release of the sterol, and diffuses into the solvent. Of note, the high level of hydration of the binding site in empty Lam4S2, i.e. after the sterol egress, is recapitulated in MD simulations of the *apo* Lam4S2 system (initiated from the sterol-free Lam4S2 structure, PDBID 6BYD) run under the same conditions as the Stage 1 simulations of the cholesterol-bound Lam4S2 described in *Figure 1* (see *Figure 2-figure supplement 4* and Methods for more details).

The sterol translocation process outlined above was sampled in its entirety in 5 out of 100 Stage 2 simulations (trajectories highlighted in red in ***Figure 2-figure supplement 5***). Another 19 trajectories in this set sampled evolution of the system from Microstate 1 to Microstate 5 (trajectories marked with a green star in ***Figure 2-figure supplement 5***), but on the time scales of these simulations, the system either did not progress further (i.e. to Microstates 6 and 7) or returned to Microstate 4 or 3 where it remained (see also below). In the remaining Stage 2 simulations the system fluctuated between Microstates 1, 2, and 3 (unmarked trajectories in ***Figure 2-figure supplement 5***).

### The side-opening to the sterol binding pocket is a key structural element of the release mechanism

The MD simulations indicate that widening of the side-opening to facilitate water penetration into the binding site (***Figure 3***) is a key step in the mechanism by which bound cholesterol leaves the protein to enter the membrane. To investigate in more detail the interplay between increased hydration of the sterol binding pocket, widening of the side-entrance to the binding cavity, and stability of cholesterol within the pocket, we analyzed the dynamics of D61 and S181 and their interactions with other residues in the binding site during the simulations. We found that D61 is engaged in electrostatic interactions with residue K89 located in the β2 strand preceding the Ω_1_ loop. Thus, the side-chain of K89 faces the entrance to the binding pocket where it interacts with the anionic side-chain of D61 (***Figure 2-figure supplement 3C, Figure 3***). This interaction is maintained in the initial stages of the translocation process (Microstates 1-4), but becomes unstable as the hydration of the sterol binding pocket reaches its highest levels after cholesterol leaves the site (note sampling of a wide range of d_61-89_ distances for Microstates 5-7 (***Figure 2-figure supplement 3C***)). These data suggest that D61, S181 and K89 together participate in stabilizing the closed conformation of the side-entrance to the binding pocket.

**Figure 3:**
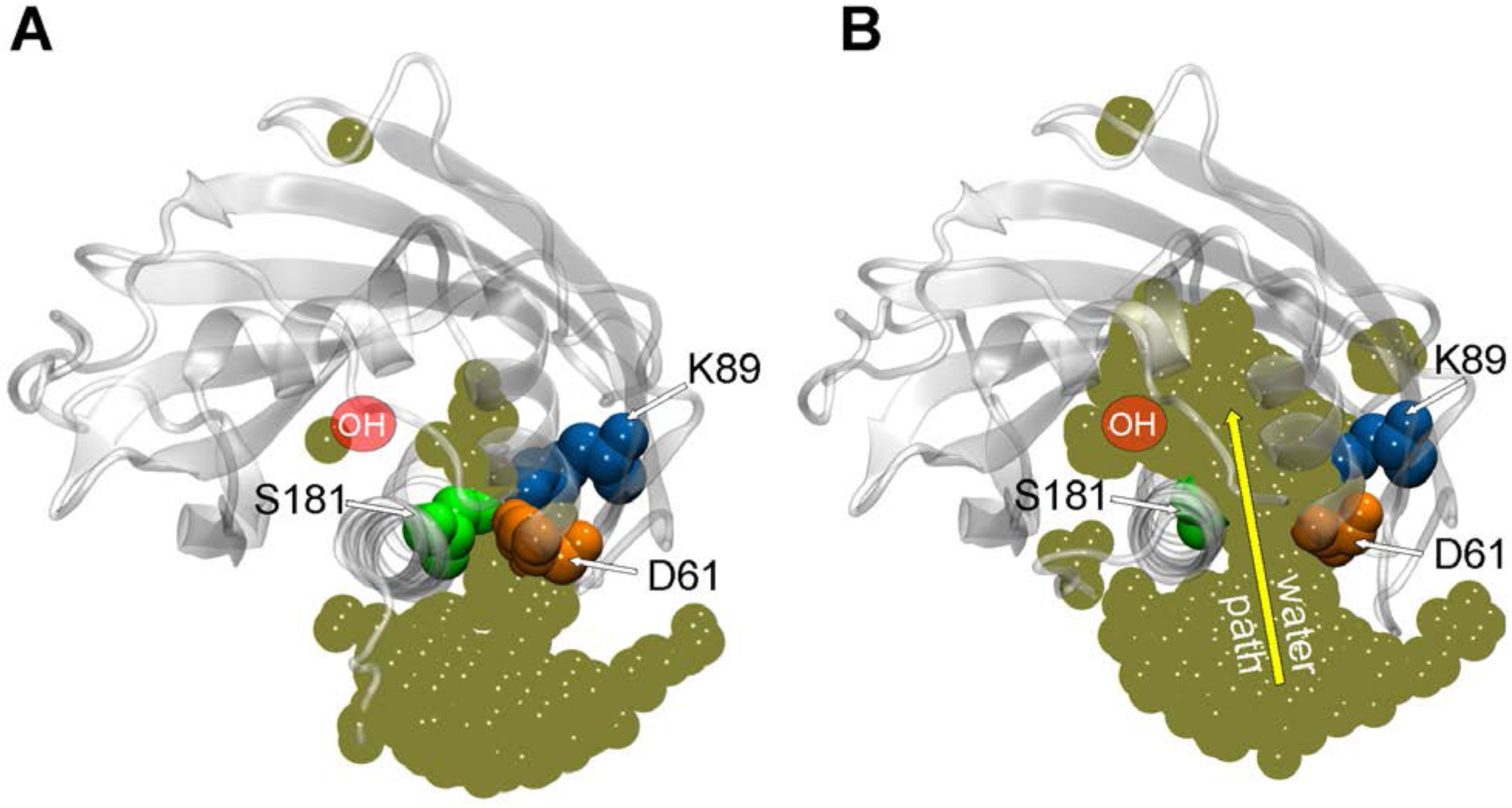
Penetration of water into the binding pocket through the side-opening is a key step in the sterol release process. (**A**-**B**) Top view of the sterol binding pocket in Lam4S2 illustrating closed (panel A) and open (panel B) conformations of the side-opening to the binding site (the protein models are representative structures from Microstates 1 and 7, respectively). In both snapshots, residues D61, K89, and S181 lining the side-opening are highlighted (in space-fill, colored and labeled). The gold spheres in panels A and B represent superposition of water oxygens in the binding site and near the side-entrance from one of the Stage 2 trajectories before (panel A) and after (panel B) the side-entrance opens. As a reference, the approximate location of the cholesterol hydroxyl group is indicated (red oval, marked OH). A water pathway to the binding pocket, formed under conditions of the open, but not closed, side-entrance is illustrated in panel B by the yellow arrow.

Based on these results and considering the position of the K89 side-chain near the protein-solvent interface, we hypothesized that replacing the polar and relatively long side-chain of K89 with a smaller hydrophobic moiety would promote widening of the side entrance, leading to destabilization of cholesterol in the binding pocket. Likewise, substituting D61 and S181 with residues with smaller size hydrophobic side-chains should have a similar destabilizing effect on bound cholesterol.

### Substitution of residues D61, S181, and K89 by Ala promotes hydration of the binding site and destabilizes bound cholesterol

To test these hypotheses, we computationally generated K89A, D61A, and S181A point-mutants of Lam4S2, and probed their dynamics using atomistic MD simulations. Specifically, we considered two snapshots taken at different time points (120 ns and 150 ns, respectively) from one of the (350 ns-long) Stage 2 trajectories of the wild type protein system in which sterol release was observed. For the wild-type protein at these time points the side-entrance to the pocket is closed (***Figure 2-figure supplement 6A***), the cholesterol molecule is stably bound (***Figure 2-figure supplement 6B***), and the level of hydration is relatively low (between 5-10 water molecules (***Figure 2-figure supplement 6B***)). We introduced the three mutations separately into these two snapshots, and - for each construct - carried out 150 ns long unbiased MD simulations in 10 replicates (1.5 µs total simulation time). Analysis of these trajectories revealed that for all the three mutants the hydration level of the sterol binding site increased rapidly, during the initial 4-5 ns of the simulations (***Figure 2-figure supplement 7A, C***; note that in the original wild type trajectory in which the mutations were introduced, reaching the same high level of hydration (>20 water molecules) required a considerably longer time (∼180 ns), see ***Figure 2-figure supplement 6B***). Furthermore, cholesterol in the trajectories for the mutant proteins was destabilized in its binding pocket (***Figure 2-figure supplement 7B, D***). Indeed, on the simulation timescales, rapid destabilization was especially notable for the K89A system in which, for all but one replicate, the sterol was unstable in its binding site (panel labeled K89A, ***Figure 2-figure supplement 7D***).

### K89A-Lam4S2 has a lower energy barrier for cholesterol release

To address the effect of the K89A mutation on cholesterol stability quantitatively, we compared the energetics of sterol release in K89A versus the wild type system using umbrella sampling MD simulations. We constructed the potential of mean force (PMF) for cholesterol release by constraining the *z*-distance between the sterol hydroxyl oxygen and the C_α_ atom of residue Q121, d_Z(chol-121)_, to different values in the range ∈ [2Å; 20 Å] along the release pathway (d_Z(chol-121)_ histogram (***Figure 2-figure supplement 3B***). The results are shown in ***Figure 4A***. For the wild type system, the PMF calculations indicate that cholesterol release requires overcoming an energy barrier of ∼6 kcal/mole, and proceeds through two major steps that were also identified in our tICA analysis of Stage 2 simulations. Thus, the PMF has a global minimum at d_Z(chol-121)_ ∼2Å corresponding to the position of cholesterol in the binding site where its polar head-group is coordinated by residue Q121 (snapshot at the top right of ***Figure 4A***), and two local minima (LM-1 and LM-2) at d_Z(chol-121)_ ∼10-14 Å and > 18Å, respectively. The global minimum represents the ensemble of states found in Microstate 1-3 (d_Z(chol-121)_ histogram, ***Figure 2-figure supplement 3B***), whereas LM-1 corresponds to the ensemble of states found in Microstate 5 (d_Z(chol-121)_ histogram, ***Figure 2-figure supplement 3B***).

**Figure 4:**
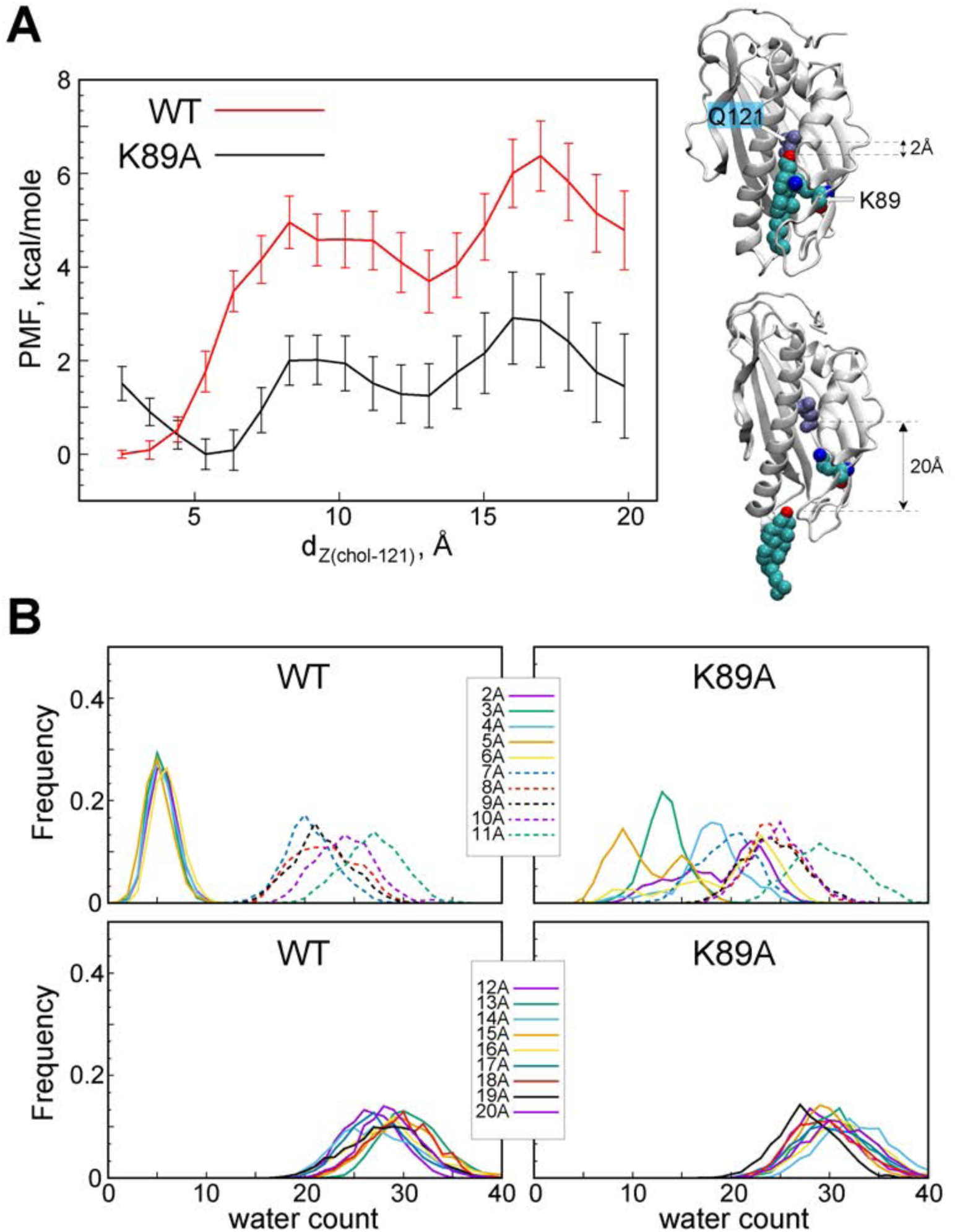
The K89A mutation reduces the energy barrier for cholesterol release. (**A**) Potential of mean force (PMF) as a function of d_Z(chol-121)_ distance for wild type (red) and K89A (black) Lam4S2 calculated from umbrella sampling MD simulations at each d_Z(chol-121)_. The structural representations on the right side of the panel illustrate locations of cholesterol corresponding to d_Z(chol-121)_ ∼2Å (top) and d_Z(chol-121)_ ∼20Å (bottom). Residues Q121 and K89 in these snapshots are also shown. (**B**) Histograms of number of water oxygens in the sterol binding site constructed from analysis of trajectories representing various windows in the range of d_Z(chol-121)_ ∈ [2Å; 20Å] from the umbrella MD simulations of the wild type (left panels) and K89A (right panels) systems.

The PMF calculations reveal that the energy barrier that separates LM-1 from the global minimum is ∼5 kcal/mole (red trace in ***Figure 4A***). This high energy cost is associated with the clear change in hydration of the sterol binding site and concomitant opening of the side-opening to the pocket (see WT profiles in ***Figure 4B***). Indeed, the water count increases and the D61-S181 interaction is destabilized when the system transitions from d_Z(chol-121)_ ∈ [2Å; 6Å] to d_Z(chol-121)_ ≥ 7Å (***Figure 4B, Figure 4-figure supplement 8A, C***). LM-2 represents the ensemble of states in which cholesterol is on the verge of exiting the protein, i.e. when it is mostly solvated by lipids and with its head-group on the level of the Ω_1_ loop (see Microstate 6 in ***Figure 2-figure supplement 3***). LM-1 and LM-2 are separated by an energy barrier of ∼2 kcal/mole. Overall, the presence of multiple minima on the PMF plot is consonant with our findings from the tICA analysis of the unbiased MD simulations described above that in some of the Stage 2 trajectories the system evolved from Microstate 1 to Microstate 5, i.e. transitioned from the global minimum to LM-1 on the PMF plot, but either did not progress further to complete sterol egress, i.e. LM-2, or returned to the conformational space of the tICA landscape characterized by relatively low hydration of the sterol binding pocket, i.e. the global energy minimum.

Remarkably, comparison of the PMF plots for the wild type and the K89A systems (***Figure 4A***) reveals that the mutation significantly lowers the barriers required for transitioning between the different energy minima. Thus, while the PMF profile for the K89A construct still has three energy minima, the energy cost to transition between the global minimum and LM-1 in this system is ∼2 kcal/mole, and between LM-2 and LM-3 is ∼1 kcal/mole, resulting in an energy barrier of only ∼3 kcal/mole for the entire release process (d_Z(chol-121)_ ∈ [2Å; 20Å]), i.e. approximately half of that determined for the wild type system. This reduction in the energy cost can be explained by a greater extent of hydration of the binding site in K89A compared to the wild type construct. Indeed, in wild type Lam4S2 the binding site remains largely dehydrated until cholesterol disengages from Q121 (d_Z(chol-121)_ ∈ [2Å; 6Å])(***Figure 4B, Figure 4-figure supplement 8A***), whereas in the K89A protein the level of solvation of the binding pocket is relatively high (> 10 water molecules) even when cholesterol is interacting with Q121 (***Figure 4B, Figure 4-figure supplement 8A***). These trends in binding site hydration are mirrored by changes in the d_61-181_ distance along the d_Z(chol-121)_ coordinate (note destabilization of D61-S181 interactions in the K89A system for small d_Z(chol-121)_ values in ***Figure 4-figure supplement 8C, D***).

Interestingly, the global minimum on the PMF profile of the K89A mutant is shifted compared to its location on the PMF plot of the wild type system from d_Z(chol-121)_ ∼ 2Å to ∼ 5Å (***Figure 4A***). We found that at the shortest d_Z(chol-121)_ distances, cholesterol-Q121 interactions in the mutant are mostly mediated by water molecules, whereas at d_Z(chol-121)_ ∼ 5Å the hydroxyl group of cholesterol is in direct contact with Q121 (see sharp peak at ∼2Å for d_Z(chol-121)_ = 5Å plot in ***Figure 4-figure supplement 8B***, note that d_Z(chol-121)_ is the Z-distance between the hydroxyl and the C_α_ of Q121). This may also explain why the water content in the cavity is skewed towards lower values for d_Z(chol-121)_ ∼ 5Å (***Figure 4B***)). Thus, for both the wild type and K89A systems, the global minimum on the PMF plot corresponds to the ensemble of states in which cholesterol is engaged in direct interactions with Q121. Overall the PMF calculations reveal that the K89A substitution lowers the energy barrier for cholesterol release from Lam4S2 into the membrane and suggests that cholesterol is consequently less stable in the binding pocket.

### Alanine substitution of residues at the side-entrance to the sterol binding pocket impacts the function of Lam4S2 in cells and *in vitro*

Our computational studies indicate that substitution of D61, K89 or S181 with alanine affects the degree of hydration of the sterol binding pocket and the stability of bound sterol, with the most significant effects seen for the K89A mutant. We tested the functionality of K89A and the other mutants using three types of experiments.

We previously showed that yeast cells lacking Ysp2/Lam2 (*lam2Δ* cells) are sensitive to the polyene antibiotic amphotericin B, and that this phenotype can be corrected by expressing a soluble GFP-Lam4S2 fusion protein (7). We verified that this was also the case for nystatin, another polyene antibiotic (***Figure 5A***, compare the first two rows in which *lam2Δ* cells are transformed with either an empty vector (row 1) or a vector for expression of GFP-Lam4S2 (row 2), and plated on media without or with different amounts of nystatin). We then tested the ability of GFP-fused Lam4S2 proteins carrying either K89A, D61A, or S181A single-point mutations (GFP-Lam4S2(K89A), GFP-Lam4S2(D61A), GFP-Lam4S2(S181A), respectively) to rescue the nystatin sensitivity of the *lam2Δ* cells. ***Figure 5A*** shows that *lam2Δ* cells expressing GFP-Lam4S2(K89A) remained nystatin-sensitive, whereas those expressing GFP-Lam4S2(D61A) or GFP-Lam4S2(S181A) became resistant to the antibiotic, similar to *lam2Δ* cells expressing wild-type protein. As all the Lam4S2 variants tested were expressed at equivalent levels (revealed by SDS-PAGE immunoblotting (***Figure 5B***)), this cell-based assay indicates that the K89A mutant has a functional deficit, whereas the D61A and S181A proteins are able to provide cells with sufficient functionality to rescue their nystatin-sensitivity phenotype.

**Figure 5:**
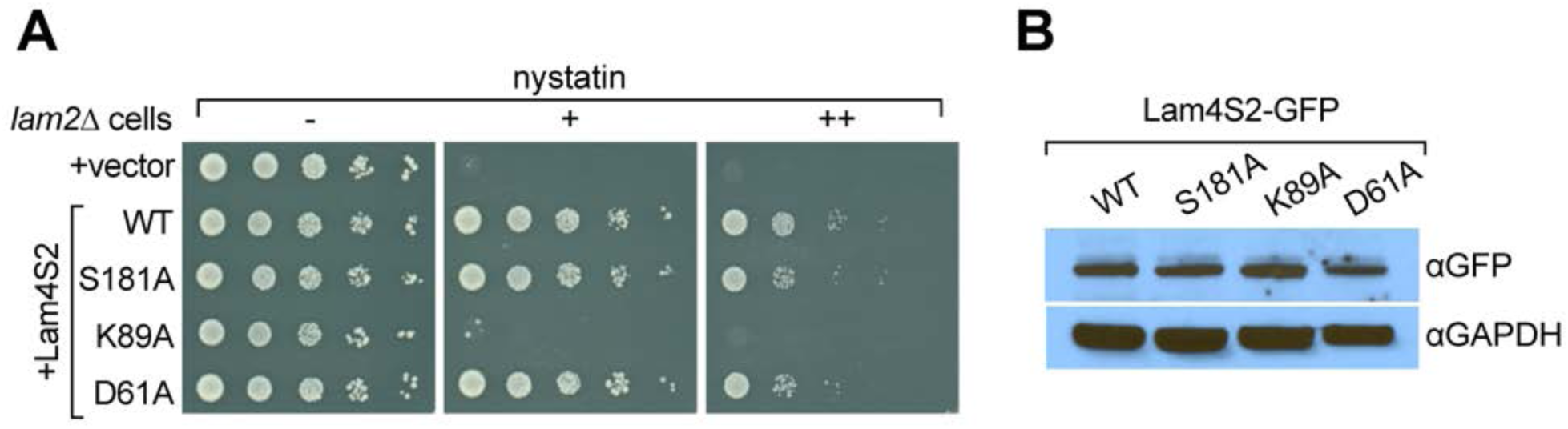
The Lam4S2(K89A) mutant does not rescue the nystatin sensitivity of *lam2Δ* cells. (**A**) Cells (*lam2Δ*) were transformed with an empty vector (top row) or with a vector for expression of GFP-Lam4S2 wild-type (WT) or mutants as indicated. Serial 10-fold dilutions were spotted onto agar plates containing defined minimal media (- uracil) lacking (-) or containing 2 μg/ml (+) or 8 μg/ml (++) nystatin. The plates were photographed after 72 h at room temperature. (**B**) Equivalent amounts of cytosol from *lam2Δ* cells expressing GFP-Lam4S2 wild-type or mutants were analyzed by SDS-PAGE and immunoblotting using anti-GFP antibodies to detect the fusion proteins and anti-GAPDH to verify equivalent loading.

To test explicitly the ability of the mutants to extract sterol from membranes and catalyze sterol exchange between populations of vesicles, we expressed His-tagged versions of the proteins in *E. coli* and purified them by affinity chromatography and size exclusion. The D61A mutant proved problematic on account of its low yield and apparent instability, and so we focused on S181A and K89A (***Figure 6A***). Similar to wild-type Lam4S2, these mutants displayed monodisperse profiles on size exclusion (***Figure 6B***) and yielded circular dichroism spectra indicative of well-folded structures (***Figure 6C***).

**Figure 6:**
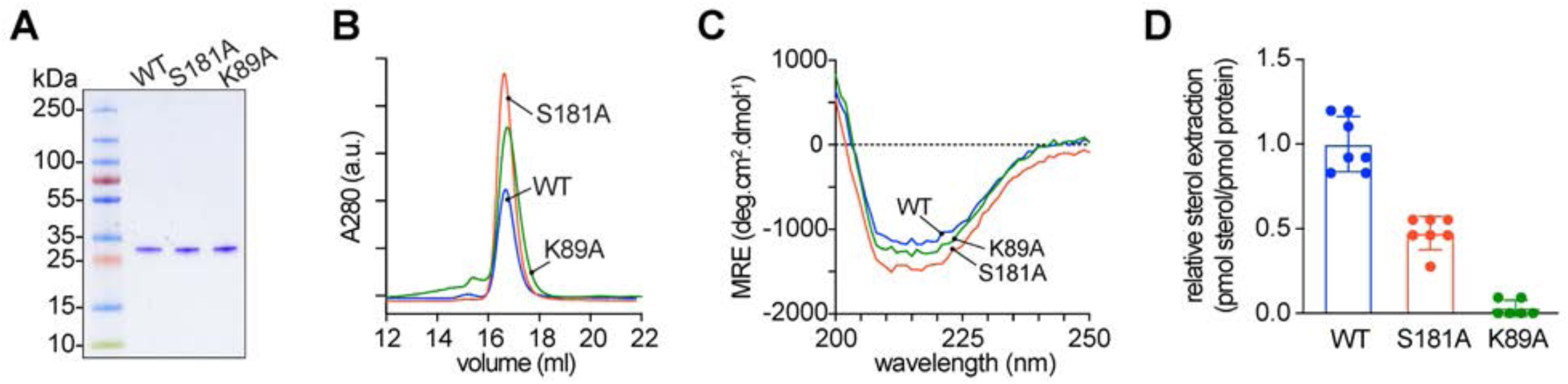
Purification and characterization of Lam4S2 mutants. (**A**) Lam4S2 wild-type and the S181A and K89A mutants were purified as His-tagged proteins via affinity chromatography and size-exclusion. The purified proteins were analyzed by SDS-PAGE (4-20% gradient gel) and Coomassie staining. (**B**) Size-exclusion analysis of purified proteins. (**C**) Circular dichroism spectra of purified proteins. Protein samples were 12 μM and the spectra shown are the average of 3 scans per sample. (**D**) Sterol extraction by purified Lam4S2 and mutants. Sucrose-loaded liposomes (DOPC:DOPE:DOPS:cholesterol, 49:23:23:5 mol %, doped with [^3^H]cholesterol) were incubated with 750 pmol of purified proteins for 1 h at room temperature. After ultracentrifugation, the radioactivity and the protein amount in the supernatant was determined, and the stoichiometry of binding was calculated. Data are represented as mean ± SEM (error bars; n = 5-7). Data are normalized to the value obtained for the wild-type protein (0.11 ± 0.02 pmol cholesterol/pmol protein (mean ± standard deviation (n=6)).

Sterol extraction assays were performed by incubating the purified proteins with large, unilamellar vesicles containing [^3^H]cholesterol, and determining the amount of radioactivity and protein in the supernatant after ultracentrifugation to pellet the vesicles. Relative to the wild-type protein, the S181A mutant extracted only ∼50% of sterol under our standard incubation conditions whereas the K89A mutant had essentially no ability to extract sterol (***Figure 6D***).

To probe sterol transfer activity of the Lam4S2 mutants, we performed *in vitro* sterol transport assays as previously described and depicted schematically in ***Figure 7A***. Donor vesicles containing fluorescent dehydroergosterol (DHE) were incubated with acceptor vesicles containing the FRET acceptor dansyl-PE. Excitation of DHE results in sensitized fluorescence emission from dansyl-PE only when the two lipids are in the same vesicle. ***Figure 7B*** (see also ***Figure 7D***) shows that under our standard conditions the wild-type protein increases the rate of DHE exchange ∼7-fold over the spontaneous rate. The S181A mutant was similar to the wild-type protein, whereas the K89A mutant had essentially no activity (***Figure 7C, D***).

**Figure 7:**
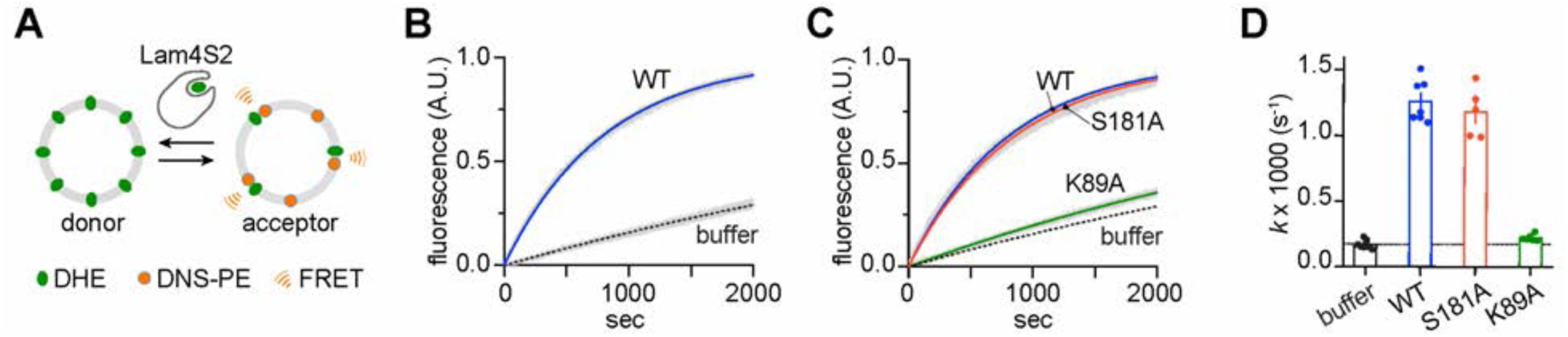
Sterol transfer activity of Lam4S2 mutants. (**A**) Schematic of the sterol transport assay. (**B**) Spontaneous sterol exchange between vesicles and transport catalyzed by wild-type Lam4S2 (0.05 μM); Traces (n=7-8) were acquired from three independent experiments and averaged. The blue and dashed lines represent monoexponential fits of the averaged data; the grey bars graphed behind the fits represent the standard error of the mean (s.e.m.). (**C**) As in panel B, except that Lam4S2 mutants were tested (n=5-7). The data fits for traces corresponding to spontaneous transport and transport catalyzed by the wild-type protein are taken from panel B and shown for comparison. (**D**) Rate constants (colored symbols) obtained from mono-exponential fits of individual traces from the experiments depicted in panels B and C. The bars show the mean and s.e.m. of the data.

Overall, the three functional tests described above indicate that the K89A mutant is compromised in sterol handling - it is unable to extract sterol from membranes and transfer it between vesicles, accounting for its inability to rescue the nystatin sensitivity of *lam2Δ* cells. These functional outcomes are in line with our computational prediction that cholesterol would be unstable in the binding site of the Lam4S2 K89A. Interestingly, the partial inability of the S181A mutant to extract cholesterol did not affect its ability to catalyze sterol exchange or rescue the nystatin sensitivity of *lam2Δ* cells.

## Discussion

Lam/GramD1 StARkin domains bind sterols specifically, admitting and exporting the sterol molecule through an aperture at the end of their long axis as suggested by inspection of crystal structures (11-14) and also seen in the MD simulations reported here. Strikingly, the sterol binding pocket in these proteins is fractured along part of its length, exposing bound sterol to solvent. The analyses presented here describe a potentially general mechanism by which sterol egress (or entry) from Lam/GramD1 StARkin domains is controlled by the concomitant entry (or egress) of water molecules via this unusual lateral fracture.

Lam4S2 engages membranes via its Ω_1_ loop and C-terminal helix, two structural regions identified previously as being functionally important in StART domains (11,12,17,18). Once membrane-bound, the protein adopts diverse conformations characterized by different extents of widening of the side-opening to the sterol binding pocket. The side-opening of the binding pocket in sterol-loaded Lam4S2 can be sealed by the polar side-chains of residues S181, D61, and K89, resulting in a low level of hydration within the cavity. In this condition, the cholesterol molecule is stably bound, with its hydroxyl group in hydrogen-bonding interactions with residue Q121. Cholesterol egress is triggered stochastically, by gradual widening of the side-opening and concomitant penetration of water into the binding site. These dynamic events destabilize cholesterol in the binding site by ∼4-5 kcal/mole, driving it from the binding site towards the membrane. The subsequent steps of the release process are enabled by repositioning of the Ω_1_ loop away from the C-terminal helix. This fully exposes the binding pocket to the membrane, i.e. widens the axial aperture, thus creating a continuous passageway to the membrane. The sequence of events by which sterol exits Lam4S2 and enters the membrane is shown in ***Figure 2-movie supplement 1***.

The overall process of cholesterol release requires overcoming an energy barrier of ∼6 kcal/mole energy barrier. This value is in a good agreement with the ∼5 kcal/mole estimate for energy barrier for sterol extraction (19) based on the assumption that intermembrane sterol transfer is rate limited by sterol pick-up/delivery processes and that the rate constant for this process can be described by simple Arrhenius relationship. Overall, the computational findings reported here reveal that the conformational state of the side-opening to the sterol binding cavity in Lam4S2 StARkin domain plays a major role in regulating the energetic stability of the sterol in the pocket.

This prediction was probed first computationally by analyzing MD trajectories of Lam4S2 in which residues that line the side-entrance to the binding site were substituted with alanine. For all three mutations (S181A, D61A and K89A) we found destabilization of cholesterol in the binding site. Using potential of mean force calculations, we found that the K89A mutation lowered the energy barrier for cholesterol release by ∼2-fold compared with wild type Lam4S2. Experimental tests confirmed that the K89A mutant was non-functional, whereas the S181A mutant was only partially compromised in its ability to bind sterol, a defect that did not appear to influence its ability to rescue the nystatin-sensitivity of *lam2Δ* cells or exchange sterols between membranes *in vitro*. Our test of the D61A mutant was limited to a cell-based assay where it performed as well as wild-type protein in rescuing the nystatin-sensitivity of *lam2Δ* cells.

Considering the functional importance of K89, and to a lesser extent S181, we examined the conservation of these residues in the Lam/GramD1 family using a previously reported structure-based sequence alignment (12). We found that the positions aligning with K89 and S181 were among the residues with the highest conservation score. Interestingly, it was noted that the side-chain of residue K910 in the S1 domain of Lam2 (Lam2S1), which aligns with K89 of Lam4S2 (note that K89 in Lam4S2 corresponds to K1031 in the full-length protein (***Table 1***)), is positioned slightly differently in the ergosterol-bound and *apo* structures (12). This led to a speculation that a path for ergosterol movement into and out of Lam2S1 could be enabled by movement of K910. Consistent with this, our study reveals that residue K89 in Lam4S2 indeed repositions when cholesterol is released from the protein. Importantly, we find that this movement is a part of larger-scale dynamic changes involving neighboring polar residues, D61 and S181, that lead to widening of the side-opening to the binding pocket.

**Table 1:** Constructs and strains.

|  |  |
| --- | --- |
| Lam4S2<sup>1</sup> | Lam4 946-1145 |
| Lam4S2(D61A)<sup>3</sup> | Lam4 946-1145 (D1003A) |
| Lam4S2(K89A)<sup>3</sup> | Lam4 946-1145 (K1031A) |
| Lam4S2(S181A)<sup>3</sup> | Lam4 946-1145 (S1123A) |
| **Yeast plasmids** |  |
| GFP only<sup>2</sup> | pRS416 (CEN *URA3*): GFP + GFP |
| GFP-Lam4S2<sup>2</sup> | pRS416 (CEN *URA3*): GFP + Lam4 946-1155 + DV<sup>4</sup> |
| GFP-Lam4S2(D61A)<sup>3</sup> | pRS416 (CEN *URA3*): GFP + Lam4 946-1155 (D1003A) + DV<sup>4</sup> |
| GFP-Lam4S2(K89A)<sup>3</sup> | pRS416 (CEN *URA3*): GFP + Lam4 946-1155 (K1031A) + DV<sup>4</sup> |
| GFP-Lam4S2(S181A)<sup>3</sup> | pRS416 (CEN *URA3*): GFP + Lam4 946-1155 (S1123A) + DV<sup>4</sup> |
| **Bacterial strain** |  |
| E. cloni EXPRESS BL21(DE3) | F– *ompT hsdSB (rB- mB-) gal dcm lon λ*(DE3 [*lacI lacUV5-T7 gene 1 ind1 sam7 nin5*]) |
| **Yeast strain** |  |
| *lam2Δ* (also called *ysp2Δ*) | *MATa his3Δ1 leu2Δ0 met15Δ0 ura3Δ0 ysp2Δ::hphNT1* |

<sup>1</sup> Described in Jentsch et al. *J. Biol. Chem.* **293**: 5522-5531 (2018)
<sup>2</sup> Described in Gatta et. al. *eLife* **4**: e07253 (2015)
<sup>3</sup> Parentheses indicate point mutations, e.g. K89A, using the Lam4S2 numbering system of Jentsch et al. (*J. Biol. Chem.* **293**: 5522-5531 (2018)); residue numbering based on the entire Lam4 sequence is provided in the right-hand column
<sup>4</sup> two amino acids (DV) appended to the end of the Lam4S2 sequence

Our computational analysis points to the key role that solvation of the sterol binding pocket plays in the process of cholesterol release. We find that water penetration destabilizes hydrogen-bonding interactions between the 3-β-OH of cholesterol and the side-chain of Q121, leading to initiation of sterol egress. While the current computations have not directly addressed the mechanism of sterol *entry* into the binding site, the PMF profile that we report here suggests that the continuous water pathway connecting the binding site to the bulk solution, as observed in our simulations of the Lam4S2 under sterol-free conditions, should play an important role in the delivery of sterol into the binding site. In this respect, we note that while in some X-ray structures of Lam/GramD1 StARkin domains the polar head-group of the bound sterol is seen in direct contact with neighboring polar residues, in the others it is engaged with the protein indirectly, through water-mediated interactions. The former mode is observed in Lam4S2, Lam2S2 and GramD1a, while the latter mode is seen in Lam2S1. Interestingly, in both Lam4S2 and Lam2S2 the head-group of the bound sterol hydrogen-bonds to the side-chain of a Gln residue (Q121 in Lam4S2). In Lam2S1, on the other hand, the position aligning with Q121 is occupied by small-size polar residue, Ser. Therefore, the head-group of the sterol does not form a direct hydrogen bond within the binding pocket of Lam2S1 but rather associates with the protein through water-mediated interactions. In GramD1a, in which the residue analogous to Q121 of Lam4S2 is also Ser, the bound sterol is seen in direct contact with another adjacent polar residue (Tyr). Taken together, the structural information highlights the importance of polar interactions for the stability of the sterol molecule in the binding site, consistent with our results demonstrating that disruption of these interactions by influx of water through the cavity side-opening leads to cholesterol release. Therefore, the molecular mechanism of sterol release that we have identified in Lam4S2 is likely to be generalizable to the other StARkin domains.

## Materials and methods

### Computational methods

#### Molecular constructs of wild type Lam4S2

The computations were based on the X-ray structures of the second StARkin domain of Lam4, Lam4S2 (PDBIDs 6BYM and 6BYD) (11). In the 6BYM structure, Lam4S2 (residue sequence 4-196 in the numbering used in Ref. (11), i.e. residue 4 corresponds to Thr-946 in native Lam4) is in complex with 25-hydroxycholesterol, which is bound in the canonical sterol binding pocket identified also in the StARkin domains of other Lam proteins (12,13). In the 6BYD model Lam4S2 (residue sequence 4-200) is in the *apo* form. For the computational studies described here, the oxysterol in the 6BYM structure was replaced by cholesterol and the molecular models of Lam4S2 in both 6BYM and 6BYD structures were completed using modeller 9v1 (20) to add respective missing residue stretches, i.e. 1-3 and 197-203 to the 6BYM structure, and 1-3 and 201-203 to the 6BYD structure.

#### Unbiased MD simulations of sterol-bound wild type Lam4S2

An all-atom model lipid membrane with the composition of “Acceptor” liposomes in sterol transport assays (11), was prepared using the CHARMM-GUI web server (21). Thus, symmetric lipid bilayer containing 70% DOPC, 15% PI, 10% DOPE, and 5% DOPS (400 lipids in total on the two leaflets) was assembled, solvated (using water/lipid number ratio of 50) and ionized with 0.1M K^+^Cl^-^ salt. This system was subjected to MD simulations for 30 ns using NAMD version 2.12 (22) and the standard multi-step equilibration protocol provided by CHARMM-GUI.

After this equilibration phase, the bilayer system was stripped of all water molecules and solution ions and the cholesterol-bound Lam4S2 domain (6BYM) was placed near the membrane surface so that the distance between any atom of the protein and any atom of the lipid molecules was ≥ 10Å (see ***Figure 1B***). The protein-membrane complex was solvated (using water/lipid number ratio of ∼145) and ionized (with 0.1M K^+^Cl^-^ salt). The resulting system contained ∼234,000 atoms in total.

The Lam4S2-membrane complex was equilibrated using a multi-step protocol (23) during which the backbone of the protein was first harmonically constrained and subsequently gradually released in three steps of 5 ns each, changing the restraining force constants from 1, to 0.5, and 0.1 kcal/ (mol Å^2^), respectively. This step was followed by 6 ns long unbiased MD simulations carried out using the NAMD 2.12 package. After this short run, the velocities of all the atoms were reset and the system was simulated with ACEMD software (24) in 10 statistically independent replicates (Stage 1 ensemble simulations), each for 320 ns, resulting in a cumulative time of 3.2 µs for Stage 1 runs.

As described in Results, Stage 1 simulations sampled events of spontaneous binding of Lam4S2 to the membrane. We randomly selected 10 frames from Stage 1 trajectories in which Lam4S2 was seen to be interacting with the lipid bilayer as in ***Figure 1E***, and initiated a new set of simulations with ACEMD (Stage 2 ensemble simulations) in which the 10 chosen structures were run in 10 statistically independent replicates each (i.e. 100 independent simulations). Each of the 100 copies were simulated for 375 ns resulting in a cumulative time of 37.5 µs for Stage 2 runs.

#### Unbiased MD simulations of *apo* wild type Lam4S2

Simulations of the *apo* wild type Lam4S2 protein (6BYD) followed the same protocol as described above for Stage 1 simulations of sterol-bound Lam4S2 with the only difference being the lipid membrane composition. Thus, in the manner identical to the sterol-bound Lam4S2, the *apo* protein was placed near the surface of the all-atom model lipid membrane (assembled with CHARMM-GUI) with the composition of “Donor” liposomes in sterol transport assays (11). This symmetric bilayer contained 31% DOPC, 23% DOPE, 23% DOPS, and 23% cholesterol (400 lipids in total on the two leaflets). As the purpose of these simulations was to quantify solvation of the empty sterol binding site, this system was only considered for Stage 1 simulations (cumulative time of 3.2 µs) and was not subjected to subsequent (Stage 2) phase.

#### Unbiased MD simulations of the mutant Lam4S2 systems

Using the FoldX server (25), three single mutations, K89A, S181A, and D79A in Lam4S2 were introduced into two separate frames of one of the Stage 2 ensemble trajectories of the wild type protein system (see Results). The resulting structures (two per mutant) were energy-minimized for 100 steps and then simulated in five independent replicates each for 150 ns using ACEMD. This resulted in 10 statistically independent MD trajectories per mutant totaling 1.5 µs.

#### Parameters and force-field for MD simulations

All the simulations performed with NAMD 2.12 implemented *all* option for rigidbonds, 2fs integration time-step, PME for electrostatics interactions (26), and were carried out in NPT ensemble under semi-isotropic pressure coupling conditions, at a temperature of 310 K. The Nose-Hoover Langevin piston algorithm (22) was used to control the target P = 1 atm pressure with the LangevinPistonPeriod set to 100 fs and LangevinPistonDecay set to 50 fs. The van der Waals interactions were calculated applying a cutoff distance of 12 Å and switching the potential from 10 Å. In addition, vdwforceswitching option was set to on.

The simulations carried out with ACEMD software implemented the PME method for electrostatic calculations, and were carried out according to the protocol developed at Acellera and implemented by us previously (24,27) with 4 fs integration time-step and the standard mass repartitioning procedure for hydrogen atoms. The computations were conducted under the NVT ensemble (at T=310 K), using the Langevin Thermostat with Langevin Damping Factor set to 0.1.

For all the simulations the CHARMM36 force field parameters for proteins, lipids, sterols, and ions (28,29) were used.

#### Umbrella sampling MD simulations of wild type and K89A Lam4S2

Biased MD simulations of cholesterol release from the wild type and the K89A mutant Lam4S2 were performed using umbrella sampling approach. The position of the translocated cholesterol was restrained to different locations along the translocation pathway (see Results) using as a collective variable the z-directional distance, d_Z(chol-121)_ (along the axis perpendicular to the membrane plane), between the cholesterol oxygen and the C_α_ atom of residue Q121 (see ***Figure 1A***). 19 windows spaced 1Å apart in the range of d_Z(chol-121)_ ∈ [2Å; 20Å] were considered and the dynamics of the sterol molecule in each window was restrained by applying a force constant of 2.5 kcal/mol · Å^2^. The rest of the parameters for the umbrella sampling runs were as follows: *width* – 2Å, and both *lowerwallconstant* and *upperwallconstant* set to 25 kcal/mol · Å^2^. Each umbrella window was simulated for 50 ns which resulted in good overlap between adjacent windows (***Figure 4-figure supplement 9A, B***).

The potential of mean force (PMF) along the collective variable was constructed with Weighted Histogram Analysis Method (WHAM) Version 2.0.9 (30). For the WHAM calculations only the last 25 ns trajectory segments of each umbrella window were used. The tolerance parameter was set to 0.0001. To estimate error bars on the PMF, for each umbrella window first decorrelation time was calculated as a time-constant from a single exponential fit to the auto-correlation vs time data (***Figure 4-figure supplement 9C, D***). The error bars were then constructed with Monte Carlo bootstrapping error analysis in the WHAM software on the decorrelated data points using *num_MC_trials* of 1000.

#### Dimensionality reduction using the time-structure based independent component analysis (tICA)

To facilitate analysis of cholesterol release process from Lam4S2 domain in the MD simulations, we performed dimensionality reduction using the tICA approach (31),(32,33),(34) as described previously (35-37). To define the tICA space we used several dynamic variables extracted from the analysis of the ensemble MD trajectories that quantify the dynamics of the cholesterol, the extent of exposure of the sterol binding site to the solvent, and the dynamics of the functionally important Ω_1_ loop. These variables include (see ***Figure 1A***): (1)-the minimum distance between the hydroxyl oxygen atom of the translocated cholesterol and residue Q121 (d_chol-121_); (2)-the root-mean-square deviation (RMSD) of the cholesterol molecule from its position in the binding site; (3)-distance between the hydroxyl oxygen of S181 and C_γ_ carbon of D61 (d_61-181_); (4)-C_α_ – C_α_ distance between residue I95 in the Ω_1_ loop and residue A169 in the C-terminal helix (d_95-169_); (5)-number of water molecules in the interior of the protein (defined as number of water oxygens found within 5Å of the side-chains of the following protein residues – 189, 185, 181, 154, 152, 136, 138, 140, 142, 123, 121, 119, 117, 102, 104, 106, 108, but farther than 5Å from the following residues – 116, 118, 109, 86, 103, 105); (6)-the number of lipid phosphate atoms with 3Å of the translocated cholesterol molecule.

Using these six CV-s as components of the data vector ***X***, the slowest reaction coordinates of a system were found as described previously (35,37,38), by constructing a time-lagged covariance matrix (TLCM): ***C***_***TL***_(*τ*)*=<****X***(*t*)***X***^*T*^(*t+τ*)*>* and the covariance matrix ***C****=<****X***(*t*)***X***^*T*^(*t*)*>*, where ***X***(*t*) is the data vector at time *t, τ* is the lag-time of the TLCM, and the symbol <…> denotes the time average. The slowest reaction coordinates are then identified by solving the generalized eigenvalue problem: ***CTLV*** = ***CVΛ***, where ***Λ*** and ***V*** are the eigenvalue and eigenvector matrices, respectively. The eigenvectors corresponding to the largest eigenvalues define the slowest reaction coordinates.

### Experimental methods

#### Lam4S2 mutants

Point mutants of Lam4S2 (D61A, K89A and S181A) were generated by PCR mutagenesis and confirmed by sequencing. The constructs and PCR primers are detailed in ***Table 1*** and ***Table 2***.

**Table 2:**
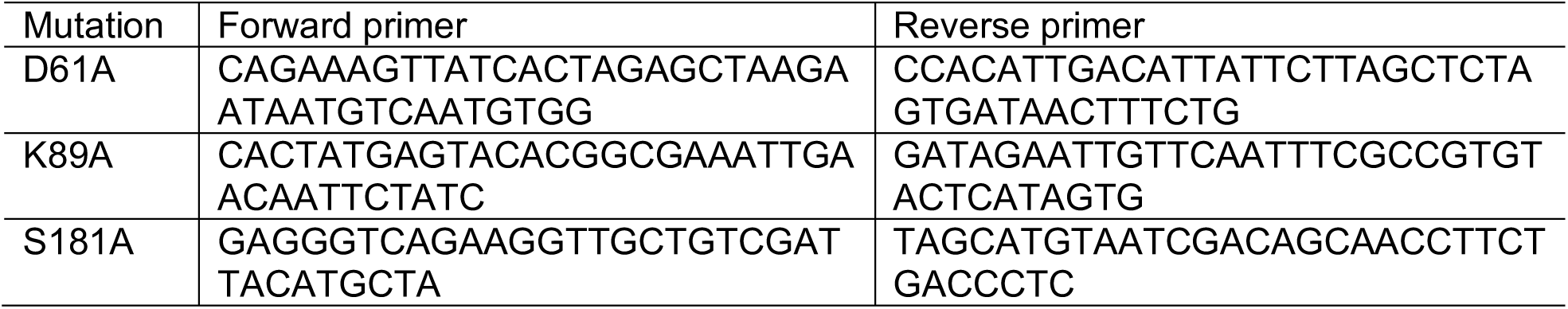
Primers used for mutagenesis.

#### Protein expression and purification

Lam4S2 and point mutants were expressed in *E. coli* as His-tagged proteins (***Table 1***), and purified by affinity chromatography on Ni-NTA resin, followed by size exclusion chromatography (SEC) using a Superdex 200 Increase 15/300GL column. The purification procedure was as previously described (11), except that the proteolysis step to remove the affinity tag was omitted and SEC was carried out in 20 mM HEPES, pH 7.5, 150 mM NaCl. The purified protein was snap frozen in small aliquots and stored at −80°C. Prior to use, aliquots were thawed and subjected to brief microcentrifugation to remove any aggregated material. Purified proteins were quantified by absorbance at 280 nm; quality control included analysis by circular dichroism (CD) as described (11), and re-analysis by SEC using buffer conditions as above.

#### Sterol transport assay

The assay (illustrated in ***Figure 7A***) was performed and analyzed as previously described (11,39) using anionic donor and acceptor liposomes (donor lipid composition: DOPC, DOPE, DOPS, DHE (31, 23, 23, and 23 mol %); acceptor lipid composition: DOPC, DOPE, liver PI, DOPS, dansyl-PE (70, 7, 15, 5, and 3 mol %). Briefly, assays were carried out at 23°C in a quartz cuvette with constant stirring using a temperature-controlled Horiba Fluoromax Plus-C spectrofluorometer; the total sample volume was 2 ml, with 0.1 mM each of donor and acceptor liposomes (final concentration, based on measurement of inorganic phosphate after acid hydrolysis of the vesicles) and 0.05 μM or 0.1 μM Lam4S2 (final concentration) in 20 mM PIPES (pH 6.8), 3 mM KCl and 10 mM NaCl (assay buffer); fluorescence was monitored for ∼2500 s using λ_ex_ = 310 nm and λ_em_ = 525 nm and a data acquisition frequency of 1 Hz. Acceptor liposomes were added to donor liposomes in the cuvette and after 60 sec, 200 µl of Lam4S2 (or Lam4S2-mutant), diluted as needed in assay buffer, was added. For control assays, 200 µl of assay buffer was added. All traces were offset corrected such that the fluorescence signal and time at the point of Lam4S2 (or buffer) addition were each set to zero. The maximum possible FRET signal was determined from assays using 0.1 µM wild-type Lam4S2 where the fluorescence readout reached a plateau value within 2000 s; traces from such assays (done in replicate) were fit to a mono-exponential function, and the plateau value obtained (FRET_max_) was used to constrain the mono-exponential fits of all other traces. Traces from different assays were compiled after data fitting by setting FRET_max_ = 1.

#### Sterol extraction assay

Sucrose-loaded liposomes were prepared as follows. Lipids (2 μmol total, of a mixture of DOPC, DOPE, DOPS, cholesterol (49, 23, 23, 5 mol %, containing a trace amount of [^3^H]cholesterol and *N*-rhodamine-DHPE) were dried in a glass screw-cap tube under a stream of nitrogen, then resuspended in 1 ml assay buffer (20 mM PIPES (pH 6.8), 3 mM KCl, 10 mM NaCl) supplemented with 250 mM sucrose, by agitating on a Vibrax orbital shaker for 30 min at 1200 rpm. The resulting suspension was subjected to 5 cycles of freeze-thaw (immersion in liquid nitrogen, followed by thawing at room temperature), before being extruded 11 times through a 200-nm membrane filter using the Avanti Mini-Extruder. After extrusion, extravesicular sucrose was removed by diluting the vesicles 4x in assay buffer and centrifuging in a Beckman TLA100.3 rotor (75,000 rpm, 1 h, 4°C). The supernatant was carefully removed from the pelleted vesicles (easily discernable because of their pink color due to rhodamine-DHPE), before resuspending the vesicles in 1 ml of assay buffer. Aliquots of the sample (5 μl) were removed at different points of preparation (after the freeze-thaw step, post-extrusion and after final resuspension) and taken for liquid scintillation counting to track lipid recovery by monitoring [^3^H]cholesterol.

The ability of Lam4S2 (wild-type and point mutants) to extract cholesterol from the vesicles was determined as follows. Liposomes (15 μl, ∼1500 pmol cholesterol) and protein (500-750 pmol, as indicated) were combined in assay buffer to a total volume of 500 μl. The mixture was incubated at room temperature for 1 h, before removing a 20 μl aliquot for liquid scintillation counting. The remainder of the sample was centrifuged in a Beckman TLA100.2 rotor (75,000 rpm, 1 h, 4°C). Most of the supernatant (350 μl) was transferred to a fresh 1.5 ml tube, while the remainder was removed immediately and discarded. The pellet was resuspended in 100 μl assay buffer containing 5% (w/v) SDS. Duplicate aliquots (50 μl each) of the supernatant were taken for liquid scintillation counting to determine the amount of extracted cholesterol. Protein in the remainder of the supernatant (250 μl) and the resuspended pellet was precipitated by adding 1.2 ml ice-cold acetone, followed by overnight incubation at −20°C. The precipitated proteins were pelleted by centrifugation, air-dried after removal of the acetone, and dissolved in SDS gel loading buffer. The relative amount of protein in the supernatant and pellet fractions was determined by SDS-PAGE, Coomassie staining and quantification of band intensity using Image J software. The data are represented as pmol cholesterol extracted / pmol protein in the supernatant.

## Acknowledgements

We thank Tim Levine and Ganiyu Alli-Balogun (Department of Cell Biology, University College London Institute of Ophthalmology) for plasmids, Trudy Ramlall (Eliezer laboratory, Weill Cornell Medical College) for help with protein purification, and Harel Weinstein for insightful comments on the manuscript. The computational work was performed using resources of the Oak Ridge Leadership Computing Facility (ALCC allocation BIP109 and Director’s Discretionary allocation) at the Oak Ridge National Laboratory, which is supported by the Office of Science of the U.S. Department of Energy under contract no. DE-AC05-00OR22725, and the computational resources of the David A. Cofrin Center for Biomedical Information in the HRH Prince Alwaleed Bin Talal Bin Abdulaziz Alsaud Institute for Computational Biomedicine at Weill Cornell Medical College.

## Additional Information

### Funding

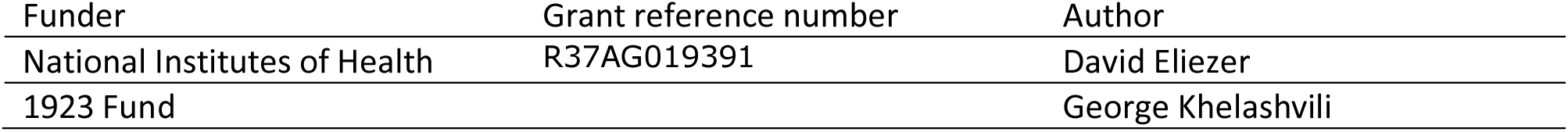

The funders had no role in study design, data collection and interpretation, or the decision to submit the work for publication.

### Author contributions

George Khelashvili, Conceptualization, Formal analysis, Investigation, Methodology, Writing— original draft, Writing—review and editing; Kalpana Pandey, Neha Chauhan, Formal analysis, Investigation, Methodology, Writing—review and editing; David Eliezer, Conceptualization, Writing—review and editing; Anant K Menon, Conceptualization, Formal analysis, Methodology, Supervision, Project administration, Writing—original draft, Writing—review and editing

## Figure Supplements

**Figure Supplement 1.**
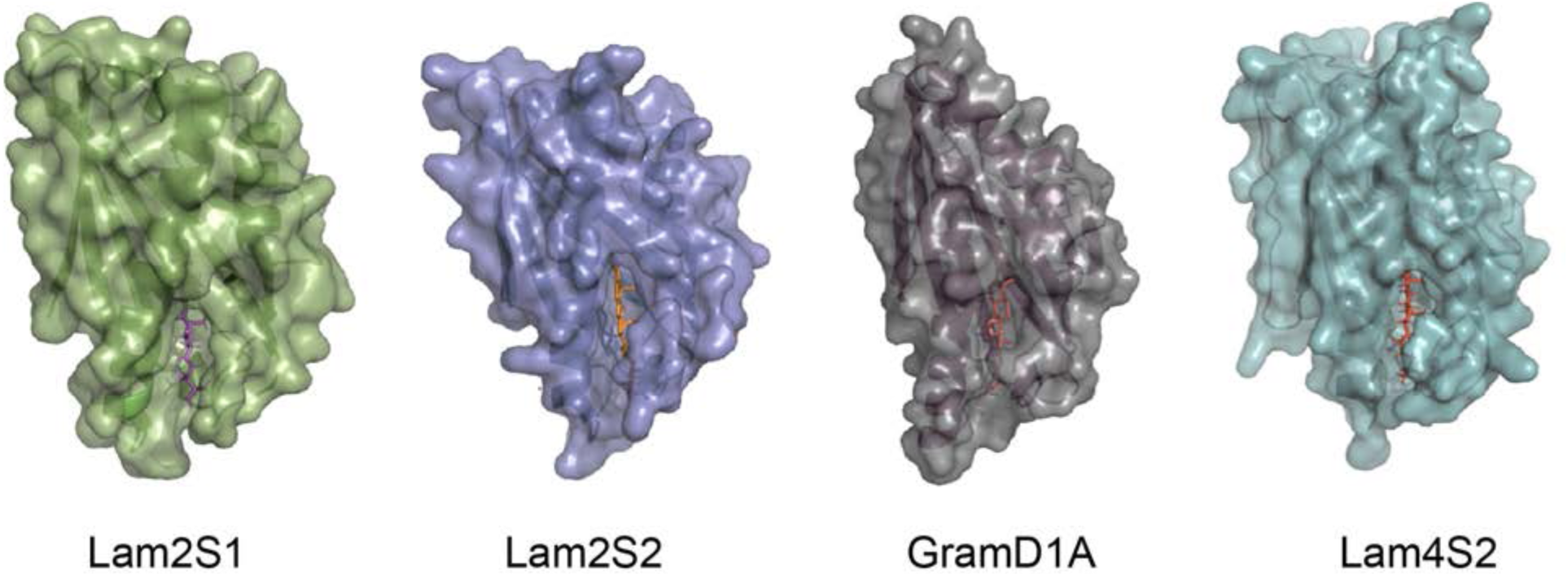
Crystal structures (surface representation) of Lam/GramD1 domains with bound sterol. The structures (PDB ID: 6CAY (Lam2S1), 5YS0 (Lam2S2), 6GQF (GramD1a), 6BYM (Lam4S2)) are oriented with their sterol entry/exit site at the bottom, and displaying the lateral fracture (side-opening) through which the bound sterol is visible from the bulk solvent.

**Figure Supplement 2:**
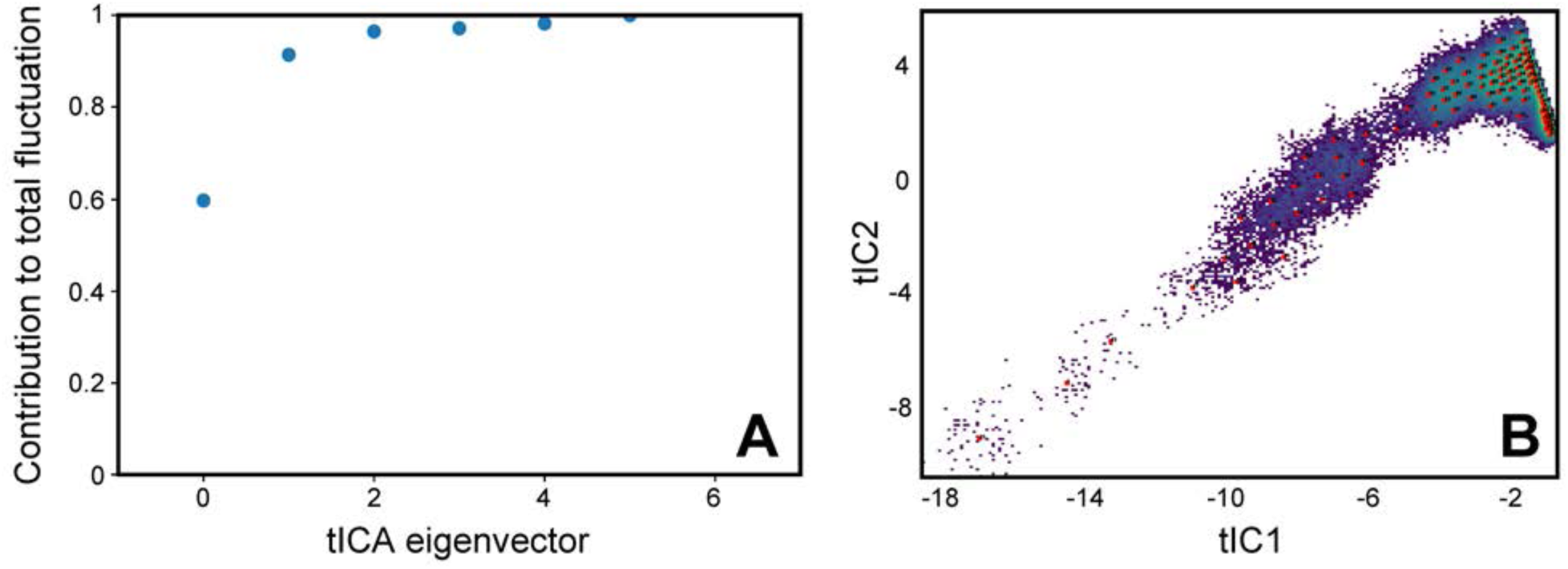
tICA analysis of Stage 2 simulations. (**A**) Contribution of each tIC vector to the total fluctuation of the system in the Stage 2 simulations. (**B**) The full set of Stage 2 trajectories mapped on the 2D landscape of the first two tICA eigenvectors (tIC 1 and tIC 2). Shown also are locations of the 100 microstates (red squares) obtained from the k-means clustering analysis of the conformational space. The lighter shades on the 2D space indicate the most populated regions (see also ***Figure 2***).

**Figure Supplement 3:**
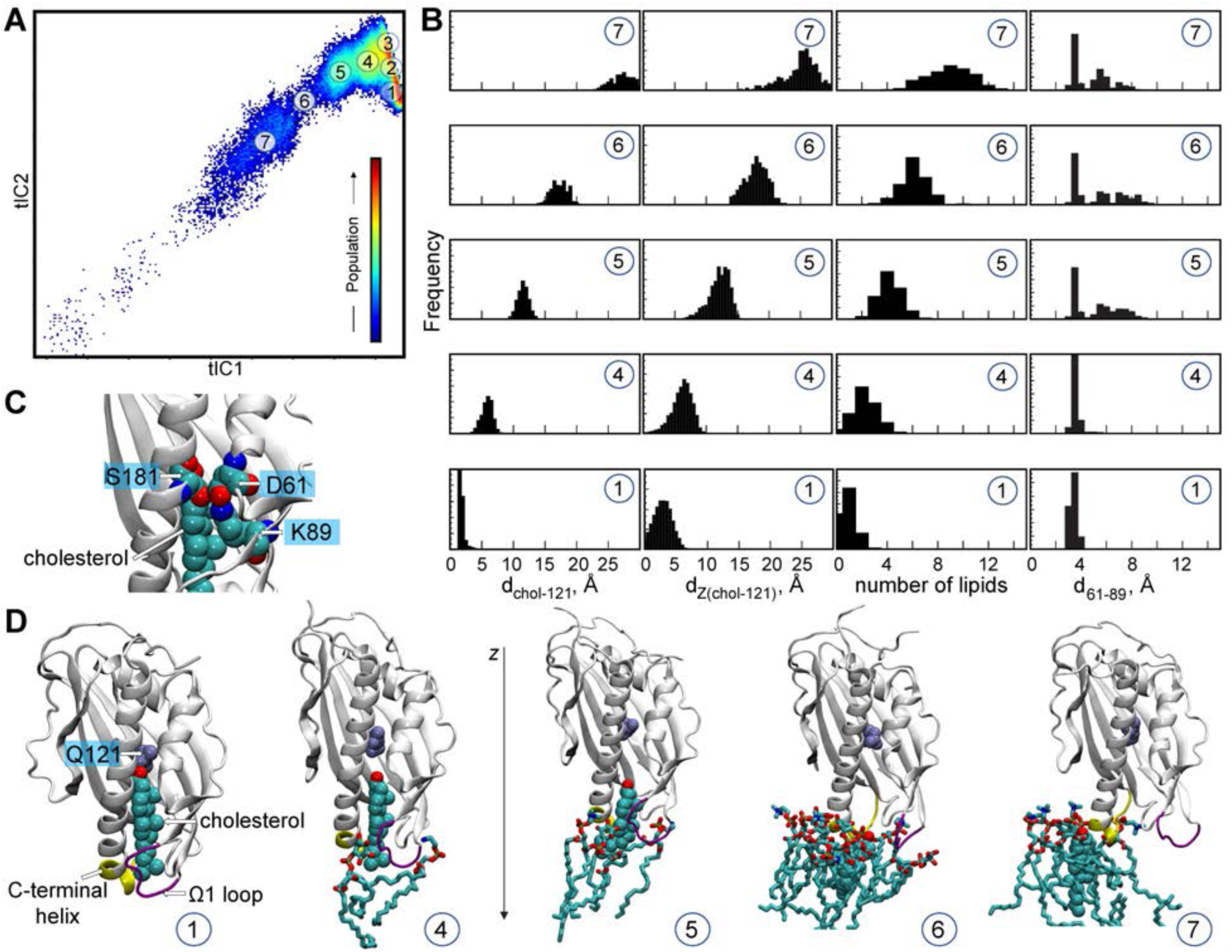
Mechanistic steps of cholesterol release from Lam4S2 revealed from tICA analysis. (**A**) 2-D landscape re-drawn from Figure 2A of the main text representing all the Stage 2 MD trajectories mapped with the tICA transformation in the space of the first two tIC vectors. The lighter shades (from red to light green to yellow) indicate the most populated regions of the 2D space (see the color bar). Microstates (see Methods) representing the most populated states in these simulations are indicated by the numbered circles (1-7) and represent various stages in the lipid translocation process. (**B**) Structural characteristics of selected microstates. The columns from left to right record the probability distributions of d_chol-121_ distance, of d_Z(chol-121)_ distance, of number of lipids within 3Å of cholesterol, and of d_61-89_ distance. (**C**) Structural snapshot zoomed at the side-entrance to the sterol binding pocket highlighting juxtaposition of D61, K89, and S181 residues (labeled). For completeness, the cholesterol is also shown in space-fill representation and is labeled. (**D**) Structural models representing microstates from panel B. These snapshots illustrate gradual immersion of the cholesterol molecule into the membrane as it exits the binding site. In these models, lipids within 3Å of cholesterol are shown in licorice, cholesterol in space-fill, and residue Q121 is colored in ice-blue and labeled. In addition, the C-terminal helix and the Ω_1_ loop are depicted in yellow and purple, respectively, and z-axis direction is highlighted.

**Figure Supplement 4:**
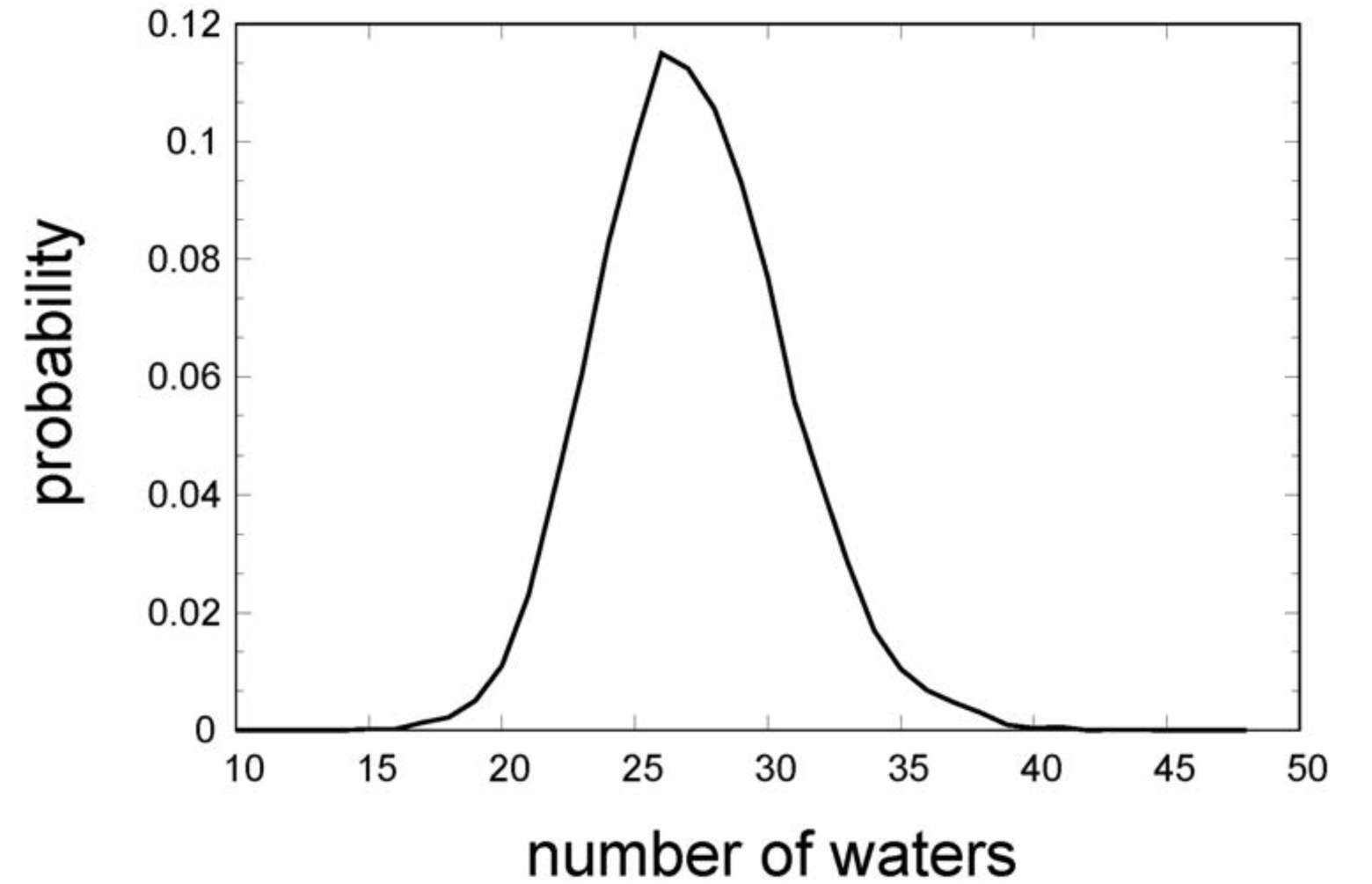
Probability distribution of the number of water oxygens in the sterol binding pocket calculated from the analysis of the Stage 1 ensemble MD simulations of *apo* Lam4S2 (based on PDBID 6BYD). The high degree of hydration of the binding pocket seen for the *apo* protein recapitulates the level of solvation observed in the MD simulations of the cholesterol-bound Lam4S2 after sterol exit (see Microstates 5-7 in ***Figure 2B***).

**Figure Supplement 5:**
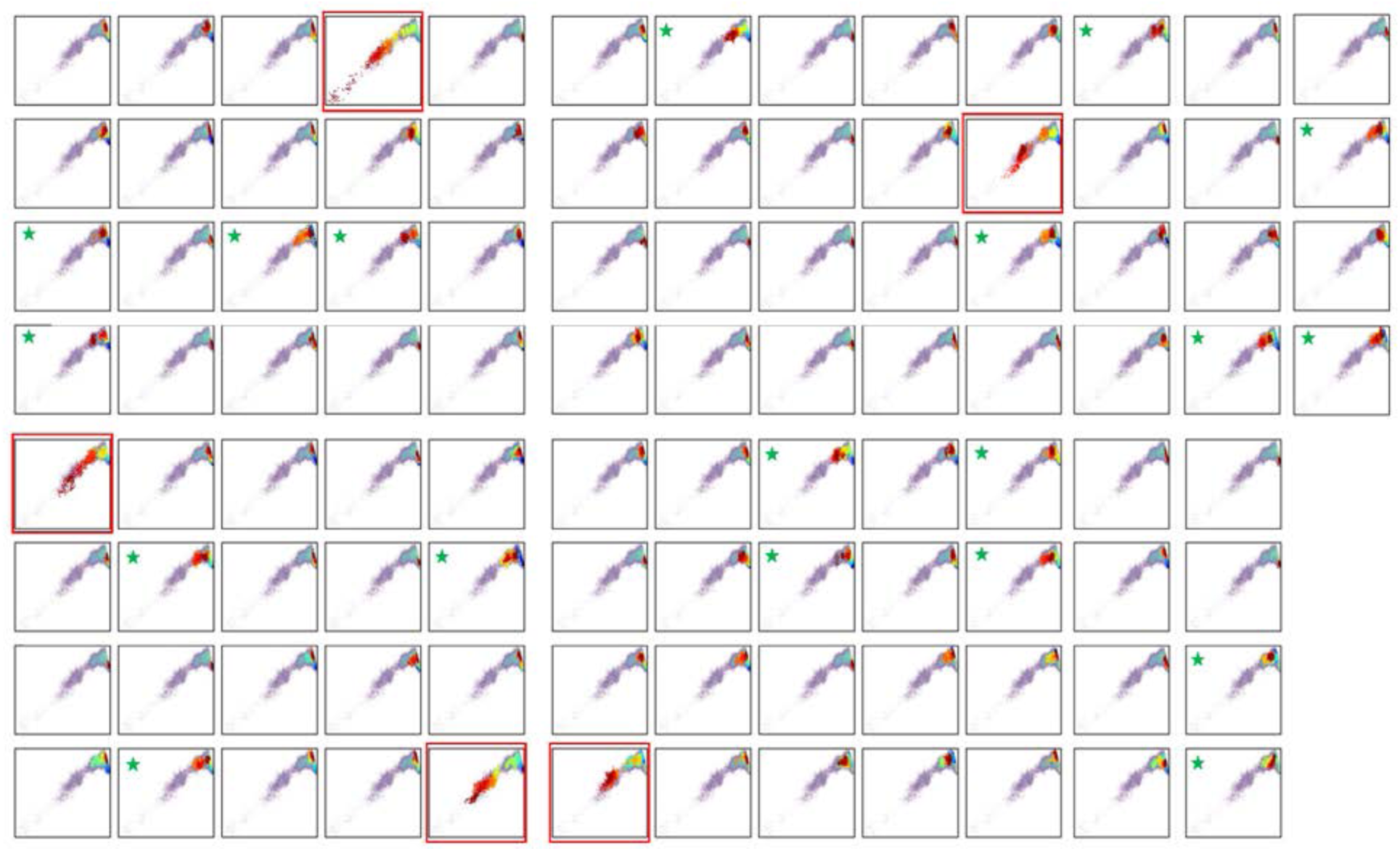
Sterol release sampled in multiple trajectories. Projection on the 2D tICA landscape from Figure 2A of each individual trajectory from Stage 2 simulations. The 2D tICA map is drawn as in Figure 2A but using smaller-size transparent dots. The colors of the larger dots indicate the time-frames in the evolution of the trajectories: darker colors (blue, cyan) represent the initial stages of the simulations, lighter colors dots (yellow, green) correspond to the middle part of the trajectories, and red shades show the last third of the trajectories. The tICA landscapes boxed in red represent simulations in which the cholesterol translocation process was sampled in its entirety. The landscapes marked with a green star are simulations in which the system evolved from Microstate 1 to Microstate 5 but did not progress further.

**Figure Supplement 6:**
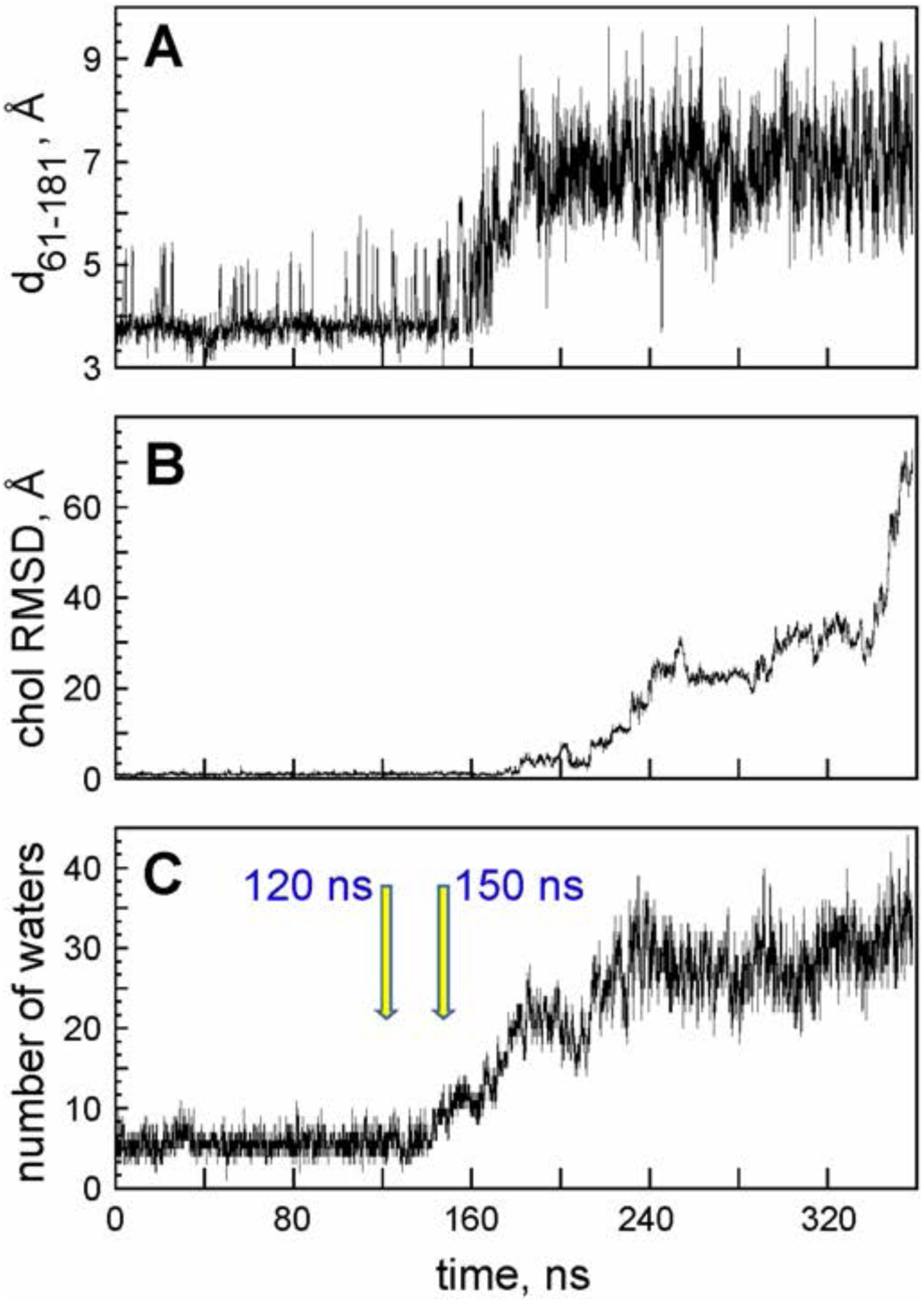
Selection of trajectory frames to initiate simulations of D61A, S181A, and K89A mutants. Time-evolution of d_61-181_ distance (**A**), cholesterol RMSD (**B**), and number of water oxygens in the sterol binding site (**C**) during one of the Stage 2 simulations in which sterol release was observed. The two trajectory frames that were selected for initiating simulations of the mutants are marked by arrows (120 ns and 150 ns time-points).

**Figure Supplement 7:**
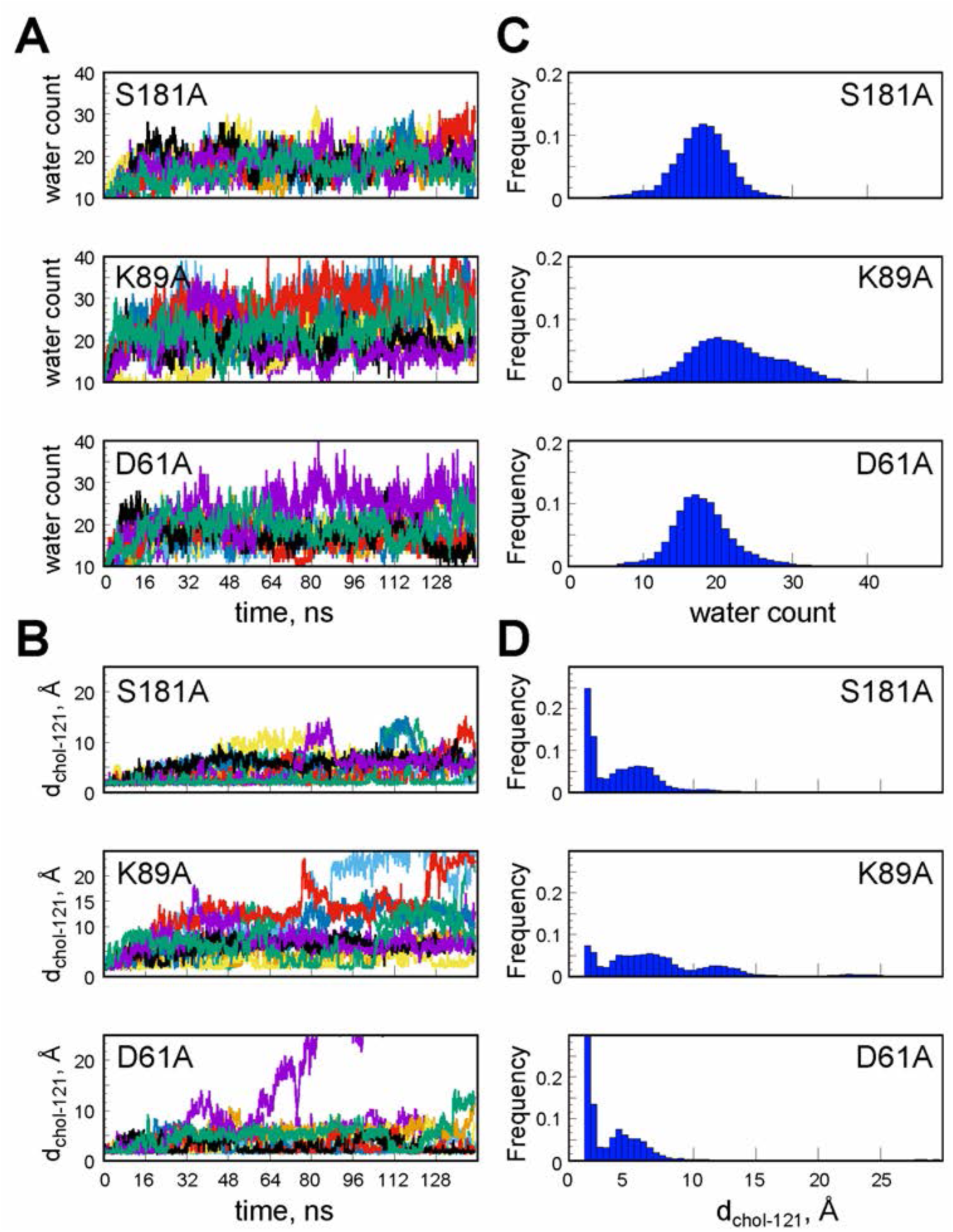
Cholesterol destabilization during unbiased ensemble MD simulations of the S181A, K89A and D61A Lam4S2 mutants. (**A**) Number of water oxygens in the sterol binding site as a function of time in simulations of S181A, K89A, and D61A Lam4S2 (top, middle, and bottom panels, respectively). The results for 10 statistically independent replicates in the corresponding ensemble of trajectories are shown in different color traces. (**B**) Minimum distance between cholesterol and Q121 residue (d_chol-121_) as a function of time in simulations of S181A, K89A, and D61A Lam4S2 (top, middle, and bottom panels, respectively). The results for 10 statistically independent replicates in the corresponding ensemble of trajectories are shown in different color traces. (**C**) Histograms of number of water oxygens corresponding to the time-traces in panel A. (**D**) Histograms of d_chol-121_ corresponding to the time-traces in panel B.

**Figure Supplement 8:**
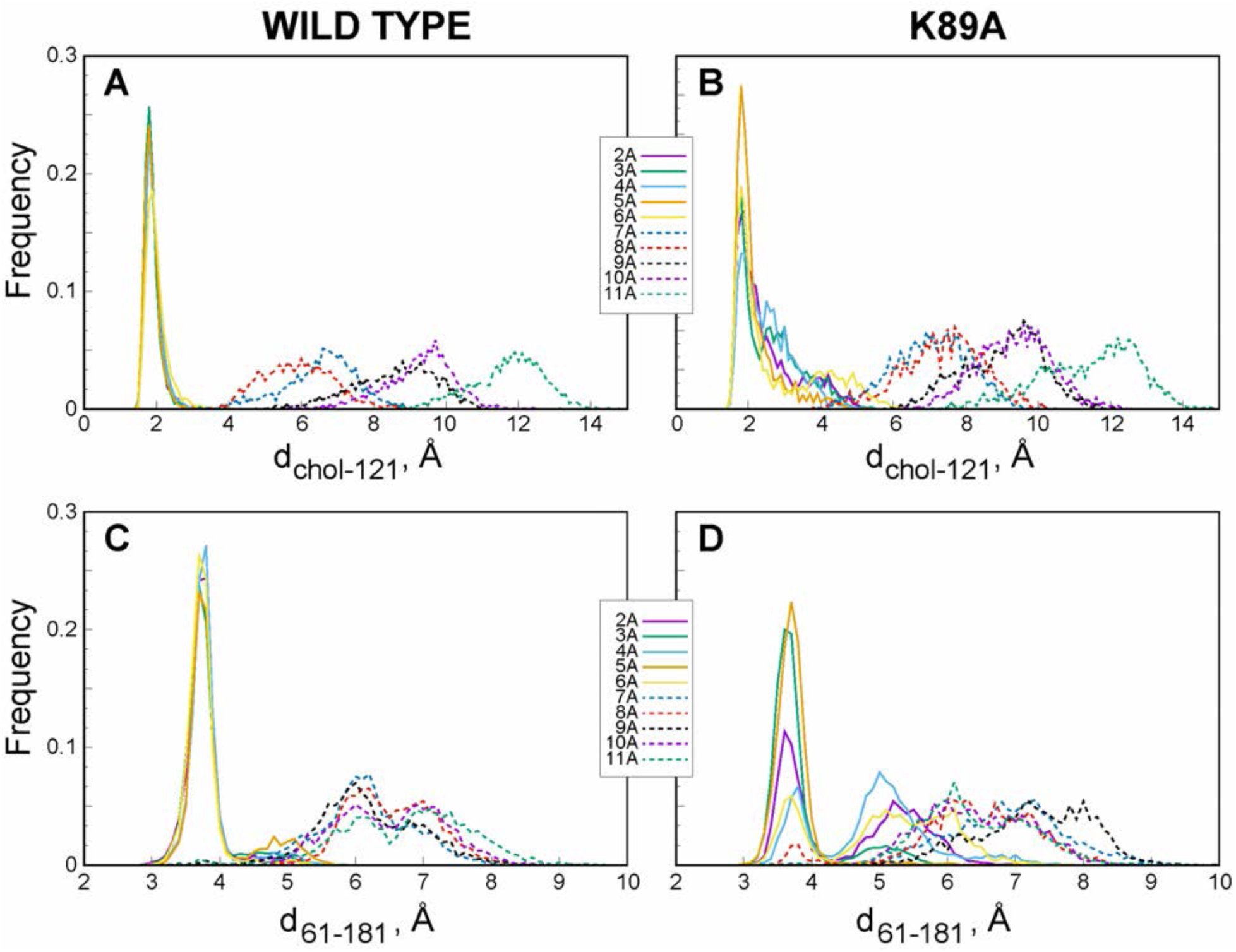
K89A mutation promotes opening of the side-entrance to the binding pocket and influx of water. Histograms of d_chol-121_ (**A**-**B**) and d_61-181_ (**C**-**D**) distances constructed from analysis of trajectories representing various windows in the range of d_Z(chol-121)_ ∈ [2Å; 20Å] from the umbrella MD simulations of the wild type (A, C) and K89A (B, D) systems.

**Figure Supplement 9:**
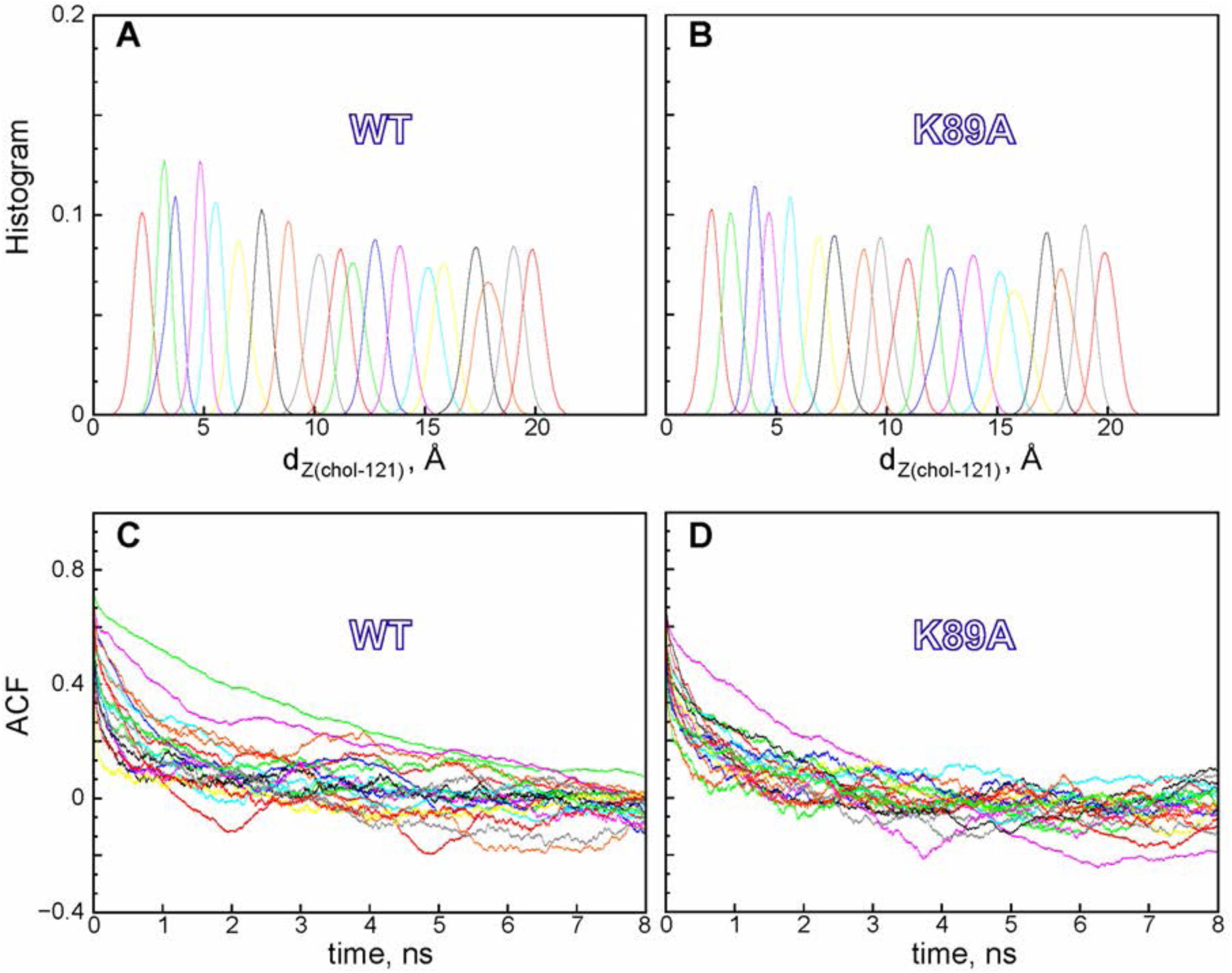
The potential of mean force calculations with WHAM. (**A**-**B**) Probability distributions of d_Z(chol-121)_ distance in different windows sampled during the umbrella MD simulations for the wild type (panel A) and the K89A mutant (panel B) systems. The data for the different umbrella windows is represented in histograms of different color. (**C**-**D**) Auto-correlation function (ACF) vs. time in different windows sampled during the umbrella MD simulations for the wild type (panel C) and the K89A mutant (panel D) systems. The color-code is the same as in panels A-B.

## Movie Supplements

**Movie Supplement 1:** Molecular dynamics trajectory of cholesterol egress from Lam4S2. The movie is based on one of the Stage 2 simulations of wild type Lam4S2. The total length of the trajectory is 350 ns. In the movie, Lam4S2 is shown in white cartoon, the cholesterol molecule is represented in ice-blue colored space-fill, S181, D61, and K89 residues are drawn in space-fill, oxygen atoms of water molecules in the sterol binding site are depicted as pink spheres, the membrane leaflet to which Lam4S2 is bound is represented by the nearby lipid phosphate atoms (golden spheres), and lipid molecules within 3Å of the cholesterol are shown in licorice representation. The rest of the simulation box is omitted. For clarity, the trajectory frames are smoothed for the movie.

## References

1. Maxfield, F. R., and van Meer, G. (2010) Cholesterol, the central lipid of mammalian cells. Curr Opin Cell Biol 22, 422–429

2. Menon, A. K. (2018) Sterol gradients in cells. Curr Opin Cell Biol 53, 37–43

3. Holthuis, J. C., and Menon, A. K. (2014) Lipid landscapes and pipelines in membrane homeostasis. Nature 510, 48–57

4. Wong, L. H., Gatta, A. T., and Levine, T. P. (2019) Lipid transfer proteins: the lipid commute via shuttles, bridges and tubes. Nat Rev Mol Cell Biol 20, 85–101

5. Wong, L. H., and Levine, T. P. (2016) Lipid transfer proteins do their thing anchored at membrane contact sites… but what is their thing? Biochem Soc Trans 44, 517–527

6. Alpy, F., and Tomasetto, C. (2005) Give lipids a START: the StAR-related lipid transfer (START) domain in mammals. J Cell Sci 118, 2791–2801

7. Gatta, A. T., Wong, L. H., Sere, Y. Y., Calderon-Norena, D. M., Cockcroft, S., Menon, A. K., and Levine, T. P. (2015) A new family of StART domain proteins at membrane contact sites has a role in ER-PM sterol transport. Elife 4

8. Elbaz-Alon, Y., Eisenberg-Bord, M., Shinder, V., Stiller, S. B., Shimoni, E., Wiedemann, N., Geiger, T., and Schuldiner, M. (2015) Lam6 Regulates the Extent of Contacts between Organelles. Cell Rep 12, 7–14

9. Murley, A., Sarsam, R. D., Toulmay, A., Yamada, J., Prinz, W. A., and Nunnari, J. (2015) Ltc1 is an ER-localized sterol transporter and a component of ER-mitochondria and ER-vacuole contacts. J Cell Biol 209, 539–548

10. Sullivan, D. P., Georgiev, A., and Menon, A. K. (2009) Tritium suicide selection identifies proteins involved in the uptake and intracellular transport of sterols in Saccharomyces cerevisiae. Eukaryot Cell 8, 161–169

11. Jentsch, J. A., Kiburu, I., Pandey, K., Timme, M., Ramlall, T., Levkau, B., Wu, J., Eliezer, D., Boudker, O., and Menon, A. K. (2018) Structural basis of sterol binding and transport by a yeast StARkin domain. J Biol Chem 293, 5522–5531

12. Horenkamp, F. A., Valverde, D. P., Nunnari, J., and Reinisch, K. M. (2018) Molecular basis for sterol transport by StART-like lipid transfer domains. EMBO J 37

13. Tong, J., Manik, M. K., and Im, Y. J. (2018) Structural basis of sterol recognition and nonvesicular transport by lipid transfer proteins anchored at membrane contact sites. Proc Natl Acad Sci U S A 115, E856–E865

14. Sandhu, J., Li, S., Fairall, L., Pfisterer, S. G., Gurnett, J. E., Xiao, X., Weston, T. A., Vashi, D., Ferrari, A., Orozco, J. L., Hartman, C. L., Strugatsky, D., Lee, S. D., He, C., Hong, C., Jiang, H., Bentolila, L. A., Gatta, A. T., Levine, T. P., Ferng, A., Lee, R., Ford, D. A., Young, S. G., Ikonen, E., Schwabe, J. W. R., and Tontonoz, P. (2018) Aster Proteins Facilitate Nonvesicular Plasma Membrane to ER Cholesterol Transport in Mammalian Cells. Cell 175, 514–529 e520

15. Quon, E., Sere, Y. Y., Chauhan, N., Johansen, J., Sullivan, D. P., Dittman, J. S., Rice, W. J., Chan, R. B., Di Paolo, G., Beh, C. T., and Menon, A. K. (2018) Endoplasmic reticulum-plasma membrane contact sites integrate sterol and phospholipid regulation. PLoS Biol 16, e2003864

16. Roelants, F. M., Chauhan, N., Muir, A., Davis, J. C., Menon, A. K., Levine, T. P., and Thorner, J. (2018) TOR complex 2-regulated protein kinase Ypk1 controls sterol distribution by inhibiting StARkin domain-containing proteins located at plasma membrane-endoplasmic reticulum contact sites. Mol Biol Cell 29, 2128–2136

17. Gatta, A. T., Sauerwein, A. C., Zhuravleva, A., Levine, T. P., and Matthews, S. (2018) Structural insights into a StART-like domain in Lam4 and its interaction with sterol ligands. Biochem Biophys Res Commun 495, 2270–2274

18. Iaea, D. B., Dikiy, I., Kiburu, I., Eliezer, D., and Maxfield, F. R. (2015) STARD4 Membrane Interactions and Sterol Binding. Biochemistry 54, 4623–4636

19. Dittman, J. S., and Menon, A. K. (2017) Speed Limits for Nonvesicular Intracellular Sterol Transport. Trends Biochem Sci 42, 90–97

20. Eswar, N., Webb, B., Marti-Renom, M. A., Madhusudhan, M. S., Eramian, D., Shen, M. Y., Pieper, U., and Sali, A. (2006) Comparative protein structure modeling using Modeller. Curr Protoc Bioinformatics Chapter 5, Unit 5 6

21. Jo, S., Lim, J. B., Klauda, J. B., and Im, W. (2009) CHARMM-GUI Membrane Builder for Mixed Bilayers and Its Application to Yeast Membranes. Biophysical Journal 97, 50–58

22. Phillips, J. C., Braun, R., Wang, W., Gumbart, J., Tajkhorshid, E., Villa, E., Chipot, C., Skeel, R. D., Kale, L., and Schulten, K. (2005) Scalable molecular dynamics with NAMD. Journal of Computational Chemistry 26, 1781–1802

23. Shi, L., Quick, M., Zhao, Y., Weinstein, H., and Javitch, J. A. (2008) The mechanism of a neurotransmitter:sodium symporter--inward release of Na+ and substrate is triggered by substrate in a second binding site. Mol Cell 30, 667–677

24. Harvey, M. J., Giupponi, G., and Fabritiis, G. D. (2009) ACEMD: Accelerating Biomolecular Dynamics in the Microsecond Time Scale. J Chem Theory Comput 5, 1632–1639

25. Schymkowitz, J., Borg, J., Stricher, F., Nys, R., Rousseau, F., and Serrano, L. (2005) The FoldX web server: an online force field. Nucleic acids research 33, W382–388

26. Essmann, U., Perera, L., Berkowitz, M. L., Darden, T., Lee, H., and Pedersen, L. G. (1995) A Smooth Particle Mesh Ewald Method. Journal of Chemical Physics 103, 8577–8593

27. Khelashvili, G., Stanley, N., Sahai, M. A., Medina, J., LeVine, M. V., Shi, L., De Fabritiis, G., and Weinstein, H. (2015) Spontaneous inward opening of the dopamine transporter is triggered by PIP2-regulated dynamics of the N-terminus. ACS Chem Neurosci 6, 1825–1837

28. Phillips, J. C., Braun, R., Wang, W., Gumbart, J., Tajkhorshid, E., Villa, E., Chipot, C., Skeel, R. D., Kale, L., and Schulten, K. (2005) Scalable molecular dynamics with NAMD. J Comput Chem 26, 1781–1802

29. Lee, J., Cheng, X., Swails, J. M., Yeom, M. S., Eastman, P. K., Lemkul, J. A., Wei, S., Buckner, J., Jeong, J. C., Qi, Y., Jo, S., Pande, V. S., Case, D. A., Brooks, C. L., 3rd, MacKerell, A. D., Jr., Klauda, J. B., and Im, W. (2016) CHARMM-GUI Input Generator for NAMD, GROMACS, AMBER, OpenMM, and CHARMM/OpenMM Simulations Using the CHARMM36 Additive Force Field. J Chem Theory Comput 12, 405–413

30. Grossfield, A. WHAM: the weighted histogram analysis method”, version 2.0.9, http://membrane.urmc.rochester.edu/wordpress/?page_id=126.

31. Molgedey, L., and Schuster, H. G. (1994) Separation of a mixture of independent signals using time delayed correlations. Physical review letters 72, 3634–3637

32. Naritomi, Y., and Fuchigami, S. (2011) Slow dynamics in protein fluctuations revealed by time-structure based independent component analysis: the case of domain motions. The Journal of chemical physics 134, 065101

33. Perez-Hernandez, G., Paul, F., Giorgino, T., De Fabritiis, G., and Noe, F. (2013) Identification of slow molecular order parameters for Markov model construction. The Journal of chemical physics 139, 015102

34. Schwantes, C. R., and Pande, V. S. (2013) Improvements in Markov State Model Construction Reveal Many Non-Native Interactions in the Folding of NTL9. J Chem Theory Comput 9, 2000–2009

35. Morra, G., Razavi, A. M., Pandey, K., Weinstein, H., Menon, A. K., and Khelashvili, G. (2018) Mechanisms of Lipid Scrambling by the G Protein-Coupled Receptor Opsin. Structure 26, 356-367.e353

36. Lee, B. C., Khelashvili, G., Falzone, M., Menon, A. K., Weinstein, H., and Accardi, A. (2018) Gating mechanism of the lipid pathway in a TMEM16 scramblase. Nature Communications 9, 3251

37. Razavi, A. M., Khelashvili, G., and Weinstein, H. (2018) How structural elements evolving from bacterial to human SLC6 transporters enabled new functional properties. BMC biology 16, 31

38. Razavi, A. M., Khelashvili, G., and Weinstein, H. (2017) A Markov State-based Quantitative Kinetic Model of Sodium Release from the Dopamine Transporter. Scientific reports 7, 40076

39. Chauhan, N., Jentsch, J. A., and Menon, A. K. (2019) Measurement of Intracellular Sterol Transport in Yeast. Methods Mol Biol 1949, 115–136

